# Astrocyte-to-microglia purinergic signaling mediates synaptic shielding and promotes neuronal activity

**DOI:** 10.64898/2026.07.05.735345

**Authors:** Koichiro Haruwaka, Juan Liu, Yue Liang, Shinya Matsumoto, Dimitrios Kleidonas, Wu Shi, Shunyi Zhao, Yong U. Liu, Anthony D. Umpierre, Zhongcong Xie, Miao Jing, Long-Jun Wu

## Abstract

How glial cells sense and regulate neuronal hypoactivity remains poorly understood. Using in vivo imaging in anesthetized, sleeping, and behaving mice, we identified a glial purinergic signaling mechanism that converts neuronal hypoactivity into local circuit regulation. Cortical hypoactivity increased spatially confined ATP release through astrocytic pannexin-1 hemichannels, which were structurally enriched near perisomatic parvalbumin-positive boutons. Local ATP release directed microglial process movement and stabilized bulbous endings through microglial P2Y12 signaling, accompanied by localized Ca²⁺ activity in microglial processes. Disruption of astrocytic pannexin-1 or microglial P2Y12 impaired bulbous-ending stabilization and abolished rebound increases in neuronal activity during emergence from anesthesia. Similar ATP release and bulbous-ending formation were observed during chemogenetic neuronal silencing or natural sleep in freely behaving mice. These findings identify astrocyte-to-microglia purinergic signaling as a mechanism that converts neuronal hypoactivity into synapse-selective microglial regulation of circuit activity.

## Introduction

Microglia are dynamic immune cells of the central nervous system that continuously survey the brain parenchyma and form activity-dependent contacts with neurons and synapses (*1–3*). In vivo imaging studies established that microglial processes rapidly extend and retract under physiological conditions, allowing microglia to monitor local tissue states with high spatial and temporal resolution (*4–7*). Recent work further showed that microglia protect neuronal function through specialized somatic purinergic junctions and limit excessive neuronal activity, indicating that microglia contribute directly to circuit homeostasis (*8, 9*).

Previous studies of activity-dependent microglial regulation have shown that neuronal hyperactivity induced microglial process extension and interaction with neural elements via adenosine triphosphate (ATP)-P2Y12 signaling (*9–13*). Interestingly, neuronal hypoactivity induced by anesthesia, whisker trimming or ocular occlusion reduces norepinephrine signaling and expands microglial surveillance territories (*14, 15*). Microglial calcium activity also increases during both neuronal hyperactivity and hypoactivity, although the physiological meaning of this response remains unclear (*16, 17*). These observations suggest that reduced neuronal activity engages active microglial process dynamics and Ca²⁺ signaling, but how these microglial responses are converted into local circuit regulation remains poorly understood. We previously showed that, during isoflurane anesthesia, microglial processes shield perisomatic inhibitory inputs onto excitatory neurons and promote neuronal activity during emergence from anesthesia (*18*). This finding revealed a functional microglial response to reduced neuronal activity, but left unresolved how neuronal hypoactivity induces spatially selective synaptic shielding by microglia toward GABAergic synapses.

Here, we show that neuronal hypoactivity engages local astrocyte-microglia signaling in the intact cortex. Hypoactive brain states induced spatially confined ATP release through astrocytic pannexin-1, which promotes microglial shielding of inhibitory synapses. These glial interactions contributed to rebound neuronal activation during emergence from anesthesia. Together, our findings identify a mechanism by which glial cells detect and locally regulate hypoactive cortical circuits.

## Results

### Anesthesia increases local cortical ATP release

Our previous results showed that microglial processes are selectively recruited to shield inhibitory synapses to promote neuronal activity during emergence from anesthesia (*18*). Consistently, motor cortical neuronal Ca²⁺ activity increased together with locomotion during emergence, and Ca²⁺ signal area correlated positively with locomotion speed (Fig. 1A to C). However, the extracellular signals that drive the intriguing selection of microglia toward inhibitory synapses during anesthesia-emergence remain unknown. We first tested whether GABA may attract microglial processes to inhibitory synapses because microglia express GABA_B_ receptors (*19, 20*). Local GABA application did not alter proximal process motility in acute cortical slices (fig. 1A to D). In contrast, ATP increased the velocity of nearby microglial processes (fig. 1E to H), consistent with previous slice imaging studies showing ATP-induced microglial process chemotaxis through P2Y receptor–dependent mechanisms (*11, 13*). These results suggest that ATP, but not GABA, could be a candidate spatial cue for microglial process remodeling.

**Fig. 1.**
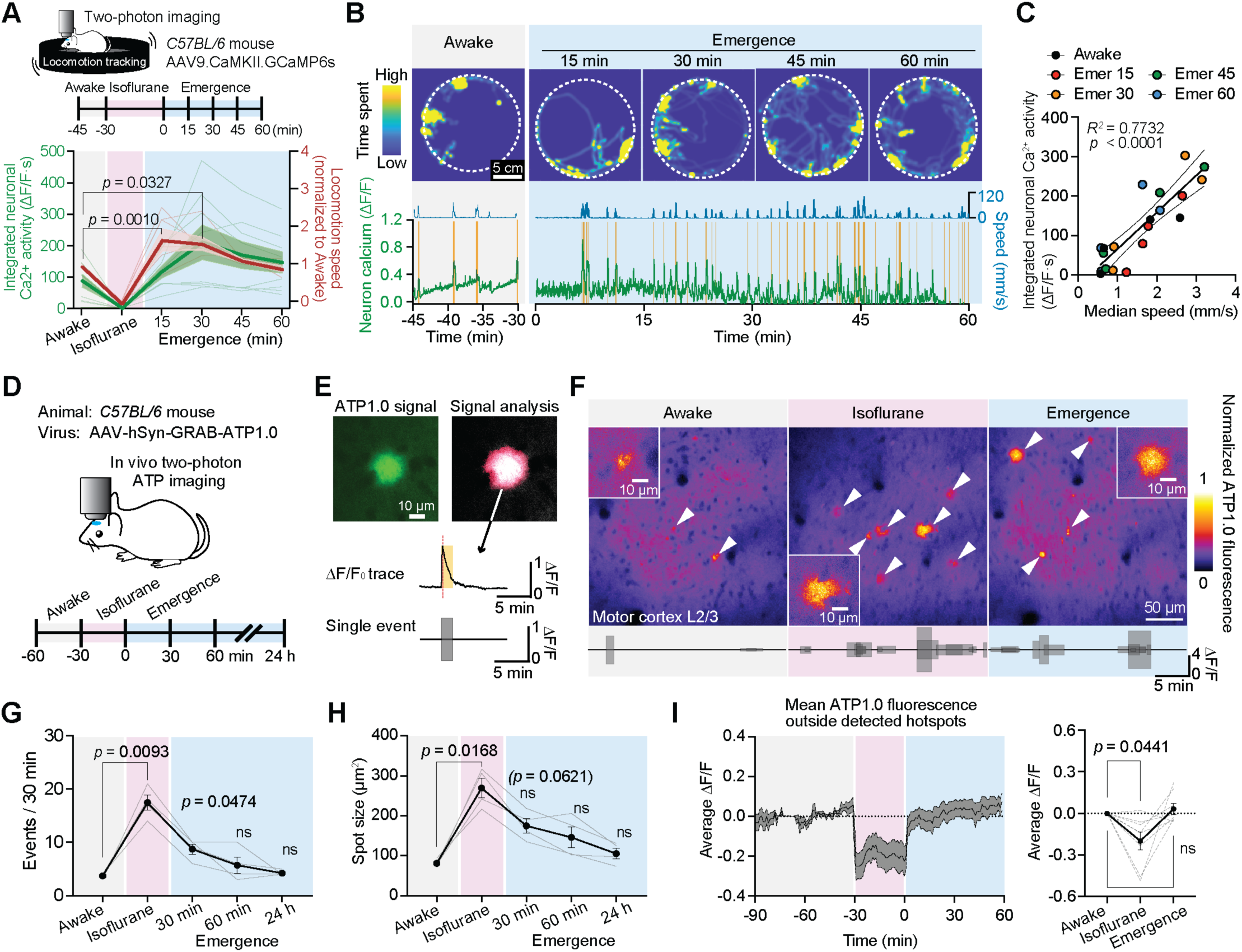
Anesthesia induces local cortical ATP release and emergence-associated rebound in neuronal activity. **(A)** Experimental design for in vivo two-photon Ca²⁺ imaging of layer 2/3 excitatory neurons in the motor cortex with simultaneous locomotion tracking, and time-course quantification of motor cortical neuronal Ca²⁺ activity and locomotion activity across awake, anesthesia, and emergence. Neuronal Ca²⁺ activity is shown as integrated ΔF/F (ΔF/F·s), and locomotion is shown as activity normalized to awake baseline. Both measurements were analyzed using one-way repeated-measures ANOVA with Dunnett’s multiple-comparisons test, n = 8 mice. **(B)** Representative locomotion occupancy maps and corresponding motor cortical neuronal Ca²⁺ activity and running speed during awake baseline and emergence. **(C)** Correlation between motor cortical neuronal Ca²⁺ activity and locomotion speed during awake baseline and emergence (linear regression analysis, n = 4 mice). **(D)** Experimental design for in vivo two-photon imaging of extracellular ATP dynamics in layer 2/3 of the motor cortex. ATP dynamics were monitored with AAV-hSyn-GRAB-ATP1.0 during awake baseline, isoflurane anesthesia, emergence from anesthesia, and 24 h after anesthesia. **(E)** Representative ATP hotspot event and schematic of ATP event detection. ATP hotspots were detected as transient GRAB-ATP fluorescence increases above a ΔF/F threshold of 0.2. The black shaded box indicates one detected ATP hotspot in the representative ΔF/F trace. **(F)** Representative pseudocolor images of normalized ATP1.0 fluorescence and corresponding single-event ΔF/F plots showing local ATP hotspot events during awake baseline, isoflurane anesthesia, and emergence. Arrowheads indicate ATP hotspots. **(G and H)** Quantification of ATP hotspot frequency (G) and ATP hotspot size (H) during awake baseline, isoflurane anesthesia, emergence, and 24 h after anesthesia (mixed-effects analysis, n = 4 mice). **(I)** Quantification of mean ATP1.0 fluorescence outside detected hotspots during awake baseline, isoflurane anesthesia, and emergence. Left, representative time course of mean non-hotspot ATP1.0 ΔF/F. Right, summary of mean ATP1.0 ΔF/F outside detected hotspots across brain states (one-way repeated-measures ANOVA with Tukey’s multiple-comparisons test, n = 8 mice). Data are shown as mean ± SEM with individual mice shown as thin lines or individual points where applicable. ns, not significant.

To examine endogenous ATP release in vivo, we used the genetically encoded G protein–coupled receptor activation–based ATP sensor (GRAB-ATP1.0) (*21*). ATP dynamics were imaged in layer 2/3 of the motor cortex across the anesthesia–emergence paradigm, with an additional 24-h recovery time point (Fig. 1D). To our surprise, we found that anesthesia increased the frequency of spatiotemporally selective hotspots, characterized as flashing fluorescence changes in ATP sensor that typically last for tens of seconds to approximately 1 min (Fig. 1E and F; Movie S1). The frequency of these ATP hotspots increased during isoflurane anesthesia, remained elevated during early emergence, and returned toward baseline by 60 min (Fig. 1G). The size of these anesthesia-induced ATP hotspots was also larger than that observed during baseline (Fig. 1H). In contrast to these spatially localized ATP release events, background ATP fluorescence outside hotspots showed a reversible decline during anesthesia and partial recovery during emergence (Fig. 1I), a pattern that may reflect reduced ambient ATP levels associated with the suppression of basal neuronal activity. Thus, localized ATP release increased throughout anesthesia and persisted into early emergence.

### Local ATP hotspots are associated with microglial bulbous-ending formation

We next simultaneously imaged ATP release (GRAB-ATP1.0), neuronal Ca²⁺ activity (CaMKII-jRGECO1a), and microglial morphology (Cx3cr1^GFP/+^ mice) in the motor cortex during isoflurane anesthesia (Fig. 2A and B; Movies S2 and S3). Similar to our previous report (*18*), microglial processes formed bulbous endings (BEs) when neuronal activity was reduced under isoflurane anesthesia. BEs were defined morphologically as terminal process enlargements that persisted for ≥5 min and had a diameter greater than 2-fold that of the adjacent process shaft. Because ATP hotspots varied in size, we classified ATP-associated BEs according to ATP hotspot area, using 100 μm² as the threshold between small and large ATP hotspots. Of all BEs detected during anesthesia, 32% were associated with large ATP hotspots (Movie S2) and 8% with small ones (Movie S3), whereas 60% were not associated with detectable ATP hotspots (Fig. 2C). Although BE-like structures could occur without detectable ATP hotspots, ATP-associated BEs persisted longer than ATP-independent BEs, with a mean lifetime of approximately 28 min (Fig. 2D). These findings suggest that local ATP release is preferentially associated with the stabilization of long-lasting BEs.

**Fig. 2.**
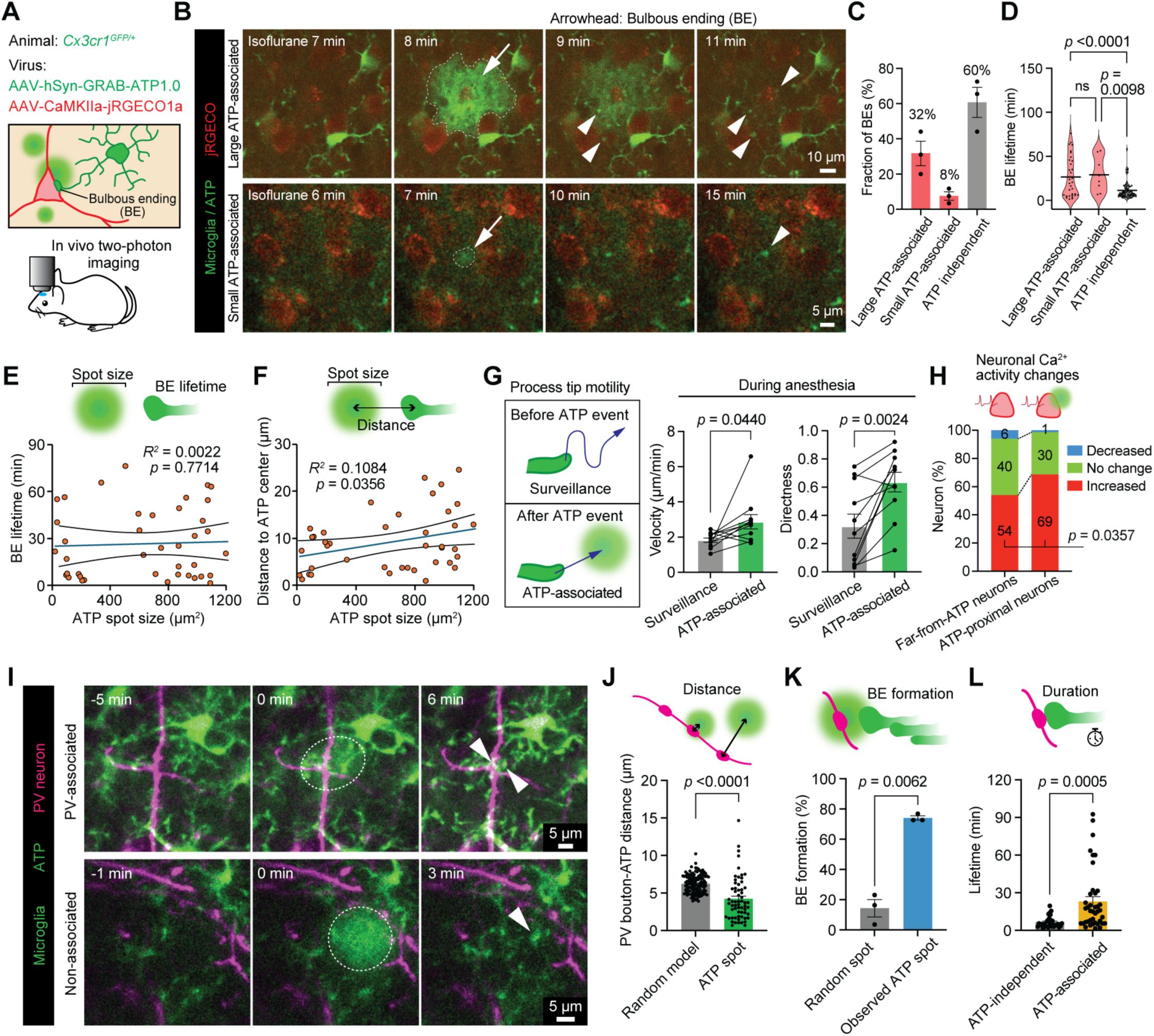
Local ATP events are associated with microglial bulbous endings near PV boutons. **(A)** Experimental design for simultaneous in vivo two-photon imaging of extracellular ATP dynamics, neuronal Ca²⁺ activity, and microglial morphology in the motor cortex during isoflurane anesthesia. ATP was monitored with GRAB-ATP1.0, neuronal activity with jRGECO1a, and microglia in Cx3cr1^GFP/+^ mice. **(B)** Representative time-lapse images showing ATP signals, microglia, and neuronal Ca²⁺ activity during isoflurane anesthesia. Arrowheads indicate bulbous endings (BEs), and the arrow indicates an ATP hotspot. **(C)** Classification of BEs according to their temporal and spatial association with ATP hotspots. BEs were categorized as large ATP-associated, small ATP-associated, or ATP-independent. ATP-associated BEs denote BEs that formed at sites of local ATP signals (n = 3 mice). **(D)** Quantification of BE lifetime for the categories shown in (C) (one-way ANOVA with Dunnett’s multiple-comparisons test; large ATP-associated, n = 32 BEs; small ATP-associated, n = 8 BEs; ATP-independent, n = 63 BEs; data collected from 5 mice). (**E and F**) Correlation analyses between ATP hotspot size and BE lifetime (E) and between ATP hotspot size and the distance from the ATP hotspot center to the associated BE (F). Linear regression; n = 41 ATP hotspots. **(G)** Quantification of microglial process tip motility during surveillance and ATP-associated movement, including velocity and directness (two-sided paired t-test, n = 11 microglia from 5 mice). **(H)** Distribution of neuronal Ca²⁺ activity changes during emergence relative to awake baseline in neurons away from ATP hotspots and in neurons close to ATP hotspots followed by BE formation. Increased and decreased activity were defined as >1.5-fold and <0.5-fold changes in Ca²⁺ signal area, respectively. Fisher’s exact test, n = 5 mice. **(I)** Representative in vivo two-photon images showing ATP hotspots, PV boutons, and microglia during isoflurane anesthesia. PV boutons were labeled with AAV-CAG-Flex-tdTomato in PV^Cre/+^::Cx3cr1^GFP/+^ mice. Arrowheads indicate BEs. **(J)** Nearest-neighbor distance between ATP hotspots and PV boutons compared with a random spatial model in which ATP hotspot positions were randomized while observed PV bouton positions were kept fixed (two-sided unpaired t-test; n = 61 ATP hotspots and 120 random points from 5 mice). **(K)** Percentage of observed ATP hotspots and matched randomized control locations associated with BE formation. Five randomized ROIs matched for ROI size, event onset time, proximity to PV boutons, and local microglial process density were generated for each observed ATP hotspot. BEs formed within 10 min of ATP event onset and persisted for ≥5 min (two-sided paired t-test, n = 3 mice). **(L)** Quantification of BE lifetime for ATP-independent and ATP-associated BEs contacting PV boutons (two-sided unpaired t-test; n = 30 ATP-independent and 40 ATP-associated BEs from 3 mice). Data are shown as mean ± SEM. ns, not significant.

We next quantified the temporal and spatial relationship between ATP hotspots and BEs. Interestingly, ATP hotspot size did not correlate with BE lifetime (Fig. 2E) but showed a positive association with the distance between the ATP hotspot center and the associated BE (Fig. 2F). Tracking of microglial process tips showed that ATP-associated movements had greater velocity and directness than surveillance movements during anesthesia (Fig. 2G). Neurons associated with ATP hotspots showed a higher fraction with increased Ca²⁺ activity during emergence, relative to awake baseline, than neurons located far from ATP hotspots (Fig. 2H).

Because ATP sensors report extracellular ATP dynamics without the cellular source, we mapped ATP hotspots relative to major cortical structures (fig. 2A to D). ATP hotspots were closest to neuronal somata across brain states, indicating that ATP events occurred near neuronal somata in the perisomatic region (fig. 2D). We further examined the spatial relationship among ATP events, microglia, and PV boutons labeled with AAV-CAG-Flex-tdTomato in PV^Cre/+^::Cx3cr1^GFP/+^ mice (Fig. 2I; Movie S4). ATP hotspots were positioned closer to PV boutons than expected from a random spatial model (Fig. 2J). Approximately 70% of ATP hotspots were followed by BE formation (Fig. 2K), and ATP-associated BEs have a longer lifetime (an average of approximately 23 min) than that of ATP-independent BEs (Fig. 2L). Together, local ATP events were associated with directed microglial process movement, BE stability, and PV bouton-proximal microglial responses.

### Astrocytic pannexin 1 is required for anesthesia-induced ATP release

Pannexin 1 (PANX1) is an ATP-permeable membrane channel implicated in astrocytic ATP release and astrocyte-to-microglia purinergic signaling under pathological brain states (*22–25*). We therefore asked whether PANX1 contributes to anesthesia-induced local ATP release using pharmacological and genetic approaches. Indeed, our results showed that PANX1 inhibition with probenecid (PBN) reduced ATP hotspot frequency during isoflurane anesthesia and emergence (Fig. 3A and B). In addition, PBN suppressed the anesthesia-induced increase in BE contacts on CaMKII-positive neuronal somata (Fig. 3C and D). Using astrocyte-specific Panx1 deletion in Aldh1l1^CreERT2^::Panx1^fl/fl^ conditional knockout (cKO) mice (Fig. 3E and F), we found that the increased ATP hotspots during isoflurane anesthesia and emergence were abolished (Fig. 3G and H). We next examined the spatial distribution of PANX1 in cortical tissue. PANX1 immunoreactivity was localized to GFAP-positive astrocytic structures near VGAT-positive inhibitory synaptic markers (Fig. 3I and fig. S3A). Nearest-neighbor analysis showed that PANX1 signals were closer to VGAT puncta than expected from a random spatial model, whereas no such proximity was observed for VGLUT1 puncta (Fig. 3J and fig. S3A, B). PANX1 proximity to VGAT or VGLUT1 puncta was unchanged by anesthesia (fig. 3C), suggesting that anesthesia did not induce acute PANX1 redistribution in astrocytes.

**Fig. 3.**
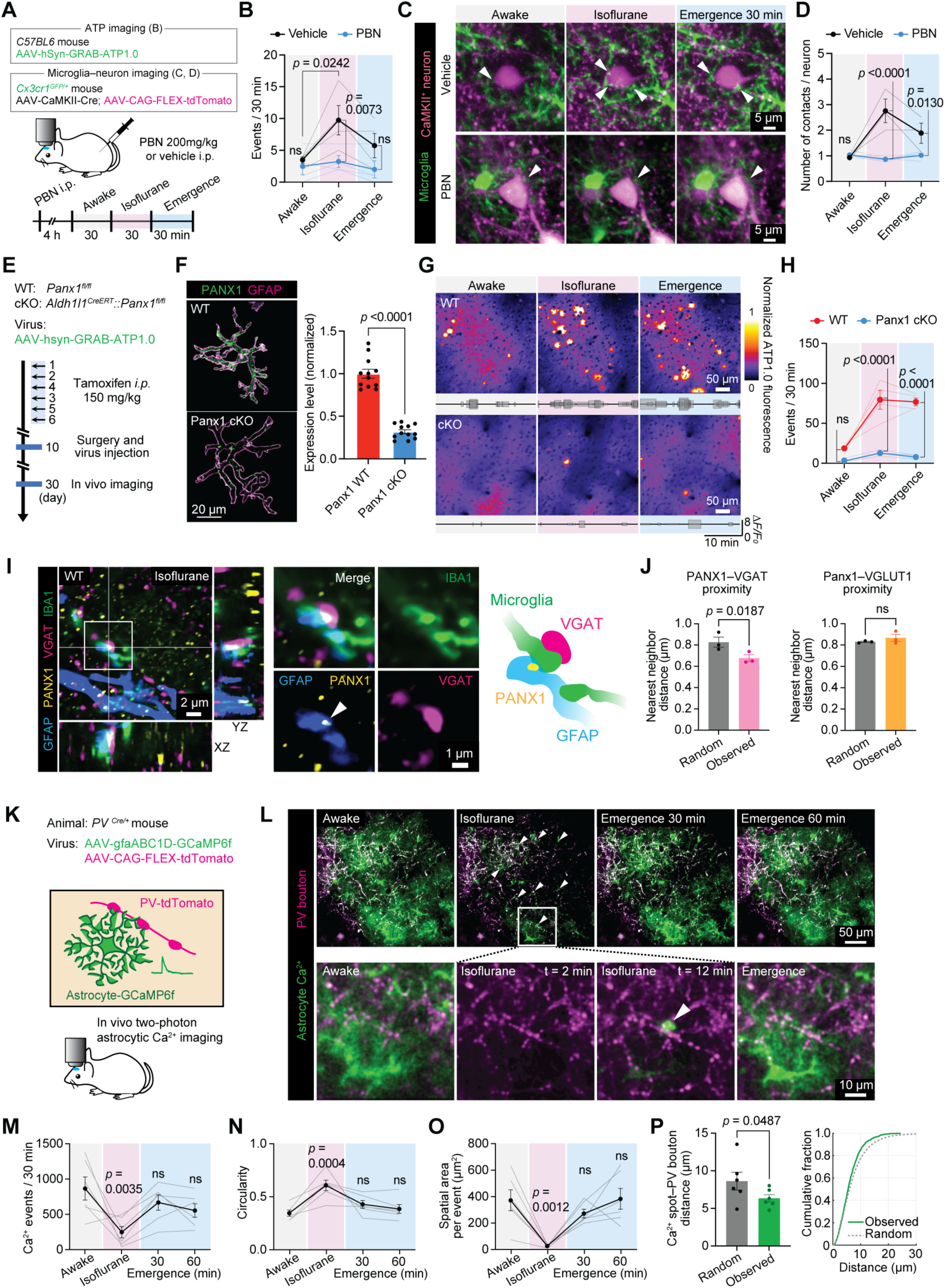
Astrocytic pannexin 1 is required for anesthesia-induced local ATP release. **(A)** Experimental design for pharmacological inhibition of pannexin 1 during in vivo imaging. C57BL/6 mice expressing GRAB-ATP1.0 in the motor cortex were used for ATP imaging and quantification, as shown in (B). For visualization of microglia–neuron interactions in (C) and quantification of BE formation in (D), Cx3cr1^GFP/+^ mice received AAV1-CaMKII-Cre together with AAV9-CAG-FLEX-tdTomato to label excitatory neurons. Mice received vehicle or probenecid (PBN) before imaging across the awake–anesthesia–emergence paradigm. **(B)** Quantification of ATP hotspot frequency in vehicle- and PBN-treated mice during awake baseline, isoflurane anesthesia, and emergence (two-way ANOVA with Tukey’s multiple-comparisons test, n = 4 mice per group). **(C)** Representative images showing BE contacts with CaMKII-positive neuronal somata in vehicle- and PBN-treated mice during awake baseline, isoflurane anesthesia, and emergence. Arrowheads indicate BE contacts. **(D)** Quantification of BE contacts per neuron in vehicle- and PBN-treated mice (two-way ANOVA with Tukey’s multiple-comparisons test; vehicle, n = 3 mice; PBN, n = 4 mice). **(E)** Experimental design for astrocyte-specific Panx1 deletion and in vivo ATP imaging in Panx1 WT and Aldh1l1^CreERT2^::Panx1^fl/fl^ conditional knockout mice. **(F)** Representative confocal images and quantification showing reduced PANX1 expression in GFAP-positive astrocytic structures in astrocytic Panx1 cKO mice compared with Panx1 WT mice (two-sided unpaired t-test; n = 8 mice per group). **(G)** Representative GRAB-ATP fluorescence images and ΔF/F traces from Panx1 WT and astrocytic Panx1 cKO mice during awake baseline, isoflurane anesthesia, and emergence. **(H)** Quantification of ATP hotspot frequency in Panx1 WT and astrocytic Panx1 cKO mice (two-way ANOVA with Tukey’s multiple-comparisons test, n = 4 mice per group). **(I)** Representative confocal images showing PANX1 immunoreactivity in GFAP-positive astrocytic structures near VGAT-positive inhibitory synaptic markers and IBA1-positive microglia during isoflurane anesthesia. Insets and schematic illustrate the spatial relationship among PANX1, GFAP, VGAT, and microglia. **(J)** Nearest-neighbor analysis of PANX1 proximity to VGAT-positive inhibitory synaptic markers and VGLUT1-positive excitatory synaptic markers compared with random spatial models (two-sided paired t-test, n = 4 mice). **(K)** Experimental design for in vivo two-photon imaging of astrocytic Ca²⁺ dynamics and PV boutons. Astrocytes were labeled with AAV-gfaABC1D-GCaMP6f and PV boutons with AAV-CAG-Flex-tdTomato in PV^Cre/+^ mice. **(L)** Representative images showing astrocytic Ca²⁺ activity and PV boutons during awake baseline, isoflurane anesthesia, and emergence. Arrowheads indicate localized astrocytic Ca²⁺ events. **(M to O)** Quantification of astrocytic Ca²⁺ event frequency (M), circularity (N), and spatial area per event (O) during awake baseline, isoflurane anesthesia, and emergence (one-way ANOVA with Dunnett’s multiple-comparisons test, n = 6 mice). **(P)** Nearest-neighbor analysis showing the distance between astrocytic Ca²⁺ events and PV boutons compared with a random spatial model (two-sided paired t-test, n = 6 mice). The cumulative distribution is shown on the right. Data are mean ± SEM. Individual points represent mice. ns, not significant.

The intracellular Ca²⁺ elevation has been reported to be a gating mechanism of PANX1 activity in astrocytes (*26*). Thus, we examined astrocytic Ca²⁺ dynamics near inhibitory synaptic sites. During isoflurane anesthesia, broad astrocytic Ca²⁺ activity was reduced (Fig. 3K to M; Movie S5), in line with previous reports (*27*). Despite the overall reduction in astrocytic Ca²⁺, the remaining events under anesthesia became more spatially confined. Individual Ca²⁺ events exhibited increased circularity and occupied a smaller spatial area (Fig. 3K to O). Interestingly, we found that these localized astrocytic Ca²⁺ events occurred closer to PV boutons than expected from a random spatial model (Fig. 3P). Together, these results indicate that anesthesia-induced ATP events require astrocytic PANX1 and occur in the presence of residual, spatially restricted astrocytic Ca²⁺ microdomain activity near PV boutons.

### Microglial P2Y12 signaling is required for ATP-associated Ca²⁺ activity and bulbous-ending stabilization

P2Y12 is a G protein–coupled purinergic receptor enriched in brain microglia that transduces chemoattractive “find-me” signals from extracellular nucleotides (*10, 11, 23*). We found that P2Y12 immunoreactivity was enriched at BEs relative to adjacent processes and somata (fig. 4A to C). Therefore, we asked whether P2Y12 signaling converts local ATP events into microglial structural and Ca²⁺ responses during anesthesia. In P2Y12 WT mice crossed with newly generated Cx3cr1-GCaMP7s-tdTomato mice (fig. 5A to D), we found that isoflurane anesthesia and emergence increased microglial Ca²⁺ event frequency, with little change in maximum amplitude (Fig. 4A to D). This state-dependent increase in event frequency was abolished in P2Y12-deficient microglia (Fig. 4C). Interestingly, we found that microglial Ca²⁺ signals were compartmentalized within BEs, with higher maximum amplitude and signal area in BEs than in adjacent processes (Fig. 4E and F; fig. 5E to G; Movie S6). Simultaneous imaging of ATP signals, microglial morphology, and microglial Ca²⁺ activity showed that ATP events in control mice were followed by BE formation and localized Ca²⁺ activity within BEs, whereas P2Y12-deficient microglia showed no detectable localized Ca²⁺ activity (Fig. 4G to I; Movies S7 and S8). BEs with localized Ca²⁺ activity persisted longer than those without detectable activity. In addition, P2Y12 deficiency shortened BE lifetime and reduced the conversion of ATP-associated swellings into stable BEs (Fig. 4J).

**Fig. 4.**
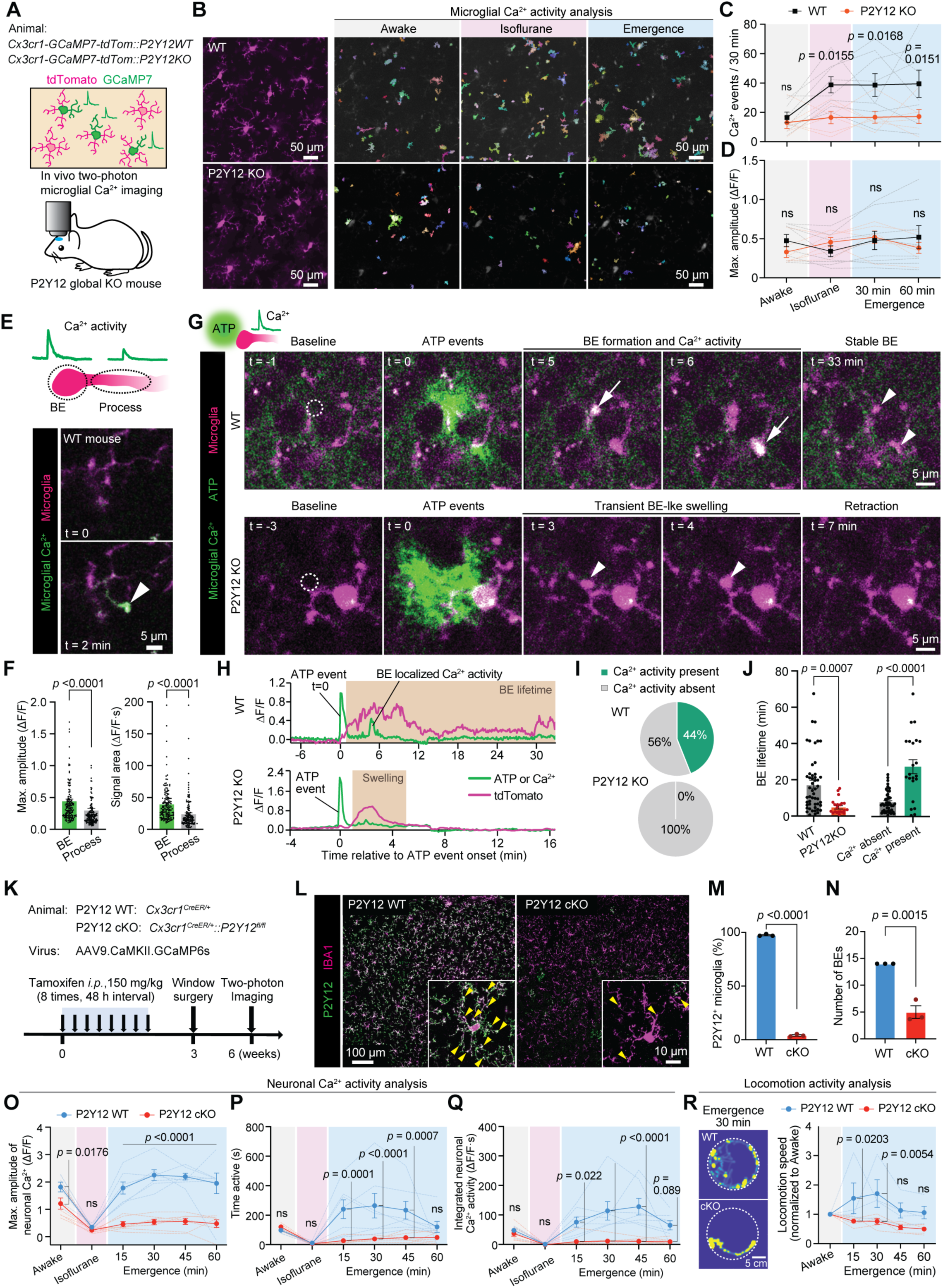
Microglial P2Y12 signaling is required for ATP-associated Ca²⁺ activity and bulbous-ending stabilization. **(A)** Experimental design for in vivo two-photon imaging of microglial Ca²⁺ activity in WT and P2Y12 KO mice. Cx3cr1-GCaMP7s-tdTomato mice were crossed with P2Y12 WT or P2Y12 KO mice. **(B)** Representative images showing microglial morphology and detected microglial Ca²⁺ events during awake baseline, isoflurane anesthesia, and emergence. Microglial morphology is shown by tdTomato fluorescence, and detected Ca²⁺ events are shown as individual event masks. **(C and D)** Quantification of microglial Ca²⁺ event frequency (C) and maximum amplitude (D) in P2Y12 WT and KO mice across brain states (two-way ANOVA with Tukey’s multiple comparisons test; WT, n = 8 mice; P2Y12 KO, n = 6 mice). **(E)** Representative images and schematic showing localized microglial Ca²⁺ activity within a BE and the adjacent process. **(F)** Event-level quantification of Ca²⁺ activity in BEs and adjacent processes in WT mice, showing maximum amplitude and signal area (two-sided paired t-test, n = 147 BE-process pairs from 4 mice). **(G)** Representative time-lapse images showing ATP hotspots, microglial morphology, and microglial Ca²⁺ activity in P2Y12 WT and KO mice during isoflurane anesthesia. Microglial Ca²⁺ events were defined as GCaMP fluorescence signals arising within microglial ROIs and completely enclosed by the tdTomato-positive microglial morphology. In WT mice, ATP events were followed by BE formation and localized microglial Ca²⁺ activity, whereas P2Y12-deficient microglia showed transient BE-like swelling followed by retraction. Arrows indicate localized microglial Ca²⁺ activity, and arrowheads indicate BEs or transient BE-like swellings. **(H)** Representative fluorescence time courses corresponding to the images in (G), showing GRAB-ATP or microglial GCaMP fluorescence together with the corresponding tdTomato fluorescence, aligned to ATP event onset. Shaded regions indicate the duration of stable BE formation in WT mice or transient BE-like swelling in P2Y12 KO mice. **(I)** Fraction of ATP-associated BEs or BE-like structures classified according to the presence or absence of localized Ca²⁺ activity in P2Y12 WT and KO mice. **(J)** Left, quantification of BE lifetime in P2Y12 WT and KO mice (two-sided unpaired t-test, n = 50 BEs from 6 WT mice and 21 BEs from 5 P2Y12 KO mice). Right, comparison of BE lifetime in WT mice with or without localized Ca²⁺ activity (two-sided unpaired t-test, n = 28 Ca²⁺ activity-absent BEs and 22 Ca²⁺ activity-present BEs from 6 WT mice). **(K)** Experimental design for microglia-specific P2Y12 deletion and in vivo imaging of neuronal Ca²⁺ activity. Cx3cr1^CreER/+^ mice were crossed with P2Y12^fl/fl^ mice, treated with tamoxifen, and subjected to cranial window surgery and two-photon imaging. **(L)** Representative confocal images showing IBA1-positive microglia and P2Y12 immunoreactivity in P2Y12 WT and microglia-specific P2Y12 conditional knockout mice. Insets show a high-magnification view of microglial processes. Arrowheads indicate P2Y12-positive microglial structures in WT mice and P2Y12-negative microglia in cKO mice. **(M)** Quantification of P2Y12-positive microglia in P2Y12 WT and microglia-specific P2Y12 cKO mice (two-sided unpaired t-test, n = 3 mice per group). **(N)** Quantification of BE number per microglial cell in P2Y12 WT and microglia-specific P2Y12 cKO mice during isoflurane anesthesia (two-sided unpaired t-test, n = 3 mice per group). **(O to Q)** Quantification of layer 2/3 neuronal Ca²⁺ activity during awake baseline, isoflurane anesthesia, and emergence in P2Y12 WT and microglia-specific P2Y12 cKO mice, including maximum amplitude (O), time active (P), and signal area (Q). Two-way ANOVA with Tukey’s multiple comparisons test; n = 7 mice per group. **(R)** Representative locomotion occupancy maps during emergence at 30 min and quantification of median locomotion speed normalized to awake baseline in P2Y12 WT and microglia-specific P2Y12 cKO mice (two-way ANOVA with Tukey’s multiple comparisons test; WT, n = 4 mice; cKO, n = 5 mice). Data are mean ± SEM. Individual points represent mice, BEs, BE-process pairs, or cells as indicated. ns, not significant.

We generated microglia-specific P2Y12 conditional knockout (cKO) mice by crossing Cx3cr1^CreER/+^ mice with P2Y12^fl/fl^ mice (*28*) (Fig. 4K). Confocal imaging confirmed loss of P2Y12 immunoreactivity in microglia of cKO mice (Fig. 4L and M). We found that P2Y12 cKO mice showed fewer BEs per microglial cell during isoflurane anesthesia than control mice (Fig. 4N). In addition, initial BE-like swellings still formed in response to ATP events in P2Y12 cKO mice, but a smaller fraction of these swellings developed into stable BEs (fig. 6A to C). We finally examined whether microglial P2Y12 signaling affects neuronal and behavioral activity during emergence from anesthesia. The emergence-associated increase in layer 2/3 neuronal Ca²⁺ responses, including maximum amplitude, time active, and signal area, was abolished in P2Y12 cKO mice (Fig. 4O to Q). Consistently, emergence-associated locomotor hyperactivity was reduced in P2Y12 cKO mice compared with controls (Fig. 4R). Together, these findings indicate that microglial P2Y12 signaling is required for persistent ATP-associated BE formation, localized microglial Ca²⁺ activity, and the rebound increase in neuronal activity during emergence from anesthesia.

### Neuronal hypoactive states trigger local ATP release and microglial bulbous-ending formation

We next tested whether chemogenetic reduction of neuronal activity could promote ATP events and microglial BE formation. To this end, we expressed Gi-DREADD in excitatory neurons together with GRAB-ATP1.0 in the motor cortex of Cx3cr1^GFP/+^ mice and monitored ATP dynamics and microglial morphology after DCZ administration (Fig. 5A and B). We found that chemogenetic suppression of excitatory neurons increased ATP hotspot frequency and spot size in the DREADD-expressing region compared with baseline (Fig. 5C and D). Spatial analysis showed that ATP hotspots were located closer to mCherry-positive DREADD-expressing neurons than expected from a random spatial model (Fig. 5E). Simultaneous imaging further revealed that microglial BEs formed near DCZ-induced ATP hotspots and DREADD-expressing neurons (Fig. 5B; Movie S9). The number of BEs per ATP hotspot increased after DCZ administration (Fig. 5F), and BE number correlated positively with ATP hotspot area after DCZ treatment (Fig. 5G). Thus, chemogenetic reduction of neuronal activity increased local ATP events and promoted microglial BE formation.

**Fig. 5.**
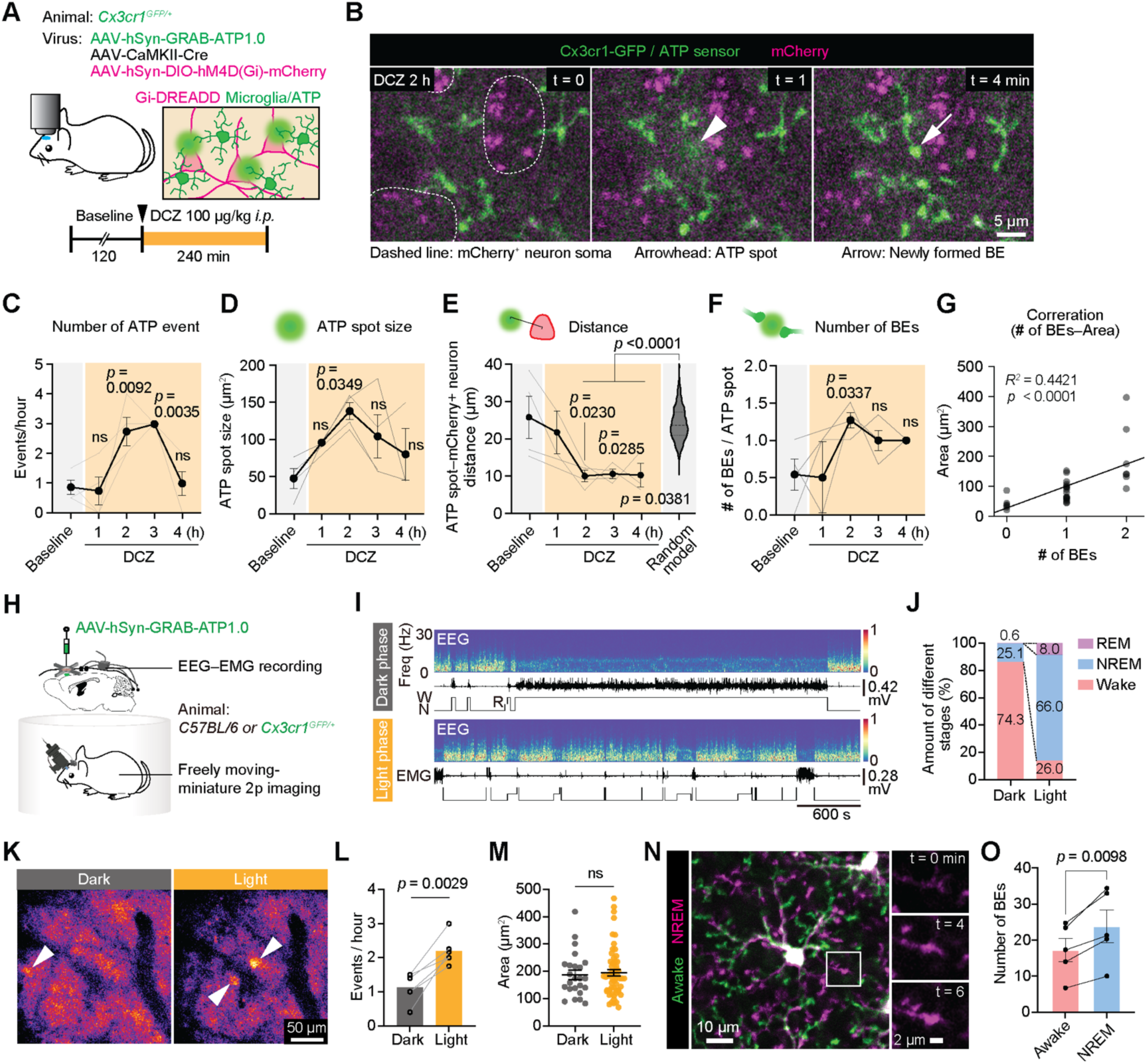
Neuronal hypoactive states trigger local ATP release and microglial bulbous ending formation. **(A)** Experimental design for chemogenetic suppression of excitatory neurons during in vivo imaging of ATP dynamics and microglial morphology. AAV-CaMKII-Cre, AAV-hSyn-DIO-hM4D(Gi)-mCherry, and AAV-hSyn-GRAB-ATP1.0 were injected into the motor cortex of Cx3cr1^GFP/+^ mice, followed by DCZ administration. **(B)** Representative time-lapse images showing ATP events and microglial morphology near mCherry-positive Gi-DREADD-expressing neurons after DCZ administration. Dashed lines indicate mCherry-positive neuronal somata. The arrowhead indicates an ATP hotspot, and the arrow indicates a newly formed BE. **(C and D)** Quantification of ATP hotspot frequency (C) and ATP hotspot size (D) before and after DCZ administration (one-way ANOVA with Dunnett’s multiple-comparisons test, n = 4 mice). **(E)** Nearest-neighbor distance between ATP hotspots and mCherry-positive Gi-DREADD-expressing neurons compared with a random spatial model (one-way ANOVA with Dunnett’s multiple-comparisons test, n = 4 mice). **(F)** Quantification of BEs per ATP hotspot before and after DCZ administration (one-way ANOVA with Dunnett’s multiple-comparisons test, n = 4 mice). **(G)** Correlation between BE number and ATP hotspot area after DCZ administration (simple linear regression; n = 30 ATP hotspots; data collected from 4 mice). **(H)** Experimental design for freely moving miniature two-photon imaging combined with EEG and EMG recordings to monitor ATP dynamics across sleep-wake states. **(I)** Representative EEG spectrograms, EMG traces, and vigilance-state classifications during dark and light phases. W, wake; N, NREM sleep; R, REM sleep. **(J)** Quantification of time spent in wake, NREM sleep, and REM sleep during dark and light phases (n = 6 mice). **(K)** Representative GRAB-ATP fluorescence images during dark and light phases. Arrowheads indicate ATP hotspots. **(L)** Quantification of ATP hotspot event frequency during dark and light phases (two-sided paired t-test, n = 6 mice). **(M)** Quantification of ATP hotspot area during dark and light phases (two-sided unpaired t-test; dark, n = 24 ATP hotspots; light, n = 57 ATP hotspots; data collected from 6 mice). **(N)** Representative images showing microglial BEs during awake and NREM sleep states. Insets show BE dynamics over time. **(O)** Quantification of BE number during awake and NREM sleep states (two-sided paired t-test, n = 5 mice). Data are shown as mean ± SEM. Individual points represent mice, ATP hotspots, or BEs as indicated. ns, not significant.

To test whether ATP events and microglial BEs also occur during physiological reductions in neuronal activity, we combined freely moving miniature two-photon imaging with EEG and EMG recordings to monitor ATP dynamics across natural sleep–wake transitions (Fig. 5H). It has been reported that sleep is associated with increased microglial Ca²⁺ activity, surveillance, and P2Y12-dependent signaling (*29, 30*). During the light phase, mice spent most of the recording period in NREM sleep, whereas wakefulness predominated during the dark phase (Fig. 5I and J). Interestingly, we found that ATP hotspots were more frequent during the light phase than during the dark phase, whereas ATP hotspot size was unchanged (Fig. 5K to M). In addition, microglial BEs were more frequent during NREM sleep than during awake periods (Fig. 5N and O). Together, these observations indicate that local ATP release and microglial BE formation are not restricted to anesthesia but also occur following chemogenetic reduction of neuronal activity or during natural sleep.

## Discussion

### Astrocytic ATP links hypoactivity to microglial engagement

How glial networks sense and regulate neuronal activity remains largely unknown. We previously reported that microglia promote neuronal activity via synaptic shielding of inhibitory synapses under anesthesia. In the present study, we further show that cortical hypoactivity induces local ATP release from astrocytes in a PANX1-dependent manner, and such ATP signaling engages microglial P2Y12 signaling to promote the BE formation near perisomatic inhibitory synapses. The local ATP release from astrocytes provides an instructive signal that links a global reduction in cortical activity to synapse-selective microglial engagement (fig. S7). Interestingly, our recent work also observed such stereotypic ATP flashes arising from astrocytes under pathological brain injury conditions, where these signals could actively present injury information to microglia and required PANX1 as the major release channel (*24*). The shared cellular and molecular architecture across these conditions suggests that PANX1-dependent astrocytic ATP signaling may serve as a common mechanism through which local brain states are communicated to microglia.

An important question is how hypoactivity triggers local astrocytic PANX1-dependent ATP release despite an overall reduction in astrocytic Ca²⁺ activity. Although PANX1 opening can be facilitated by intracellular Ca²⁺, the channel is also regulated by a broad range of mechanisms, including membrane potential, ionic composition, and post-translational modifications. How astrocytic PANX1 is gated under physiological conditions remains largely unclear. Astrocytes exhibit highly localized Ca²⁺ microdomain activity within their fine processes, which can occur independently of coordinated cell-wide Ca²⁺ signals (*31*). Our results indicate that anesthesia does not homogeneously suppress astrocytic Ca²⁺ activity; rather, small, spatially confined Ca²⁺ events persisted near inhibitory synaptic regions, where astrocytic PANX1 is also enriched. Such local Ca²⁺ events may engage spatially restricted signaling pathways for PANX1 activity. Future studies will be required to determine how neuronal hypoactivity shapes local astrocytic Ca²⁺ signaling and how these signals are coupled to PANX1 gating and ATP release. More broadly, recent studies have established astrocyte-microglia communication through a variety of signaling, including Wnts, IL-3, and complement C3, in brain development and diseases (*32–34*). Our findings add spatially confined astrocytic ATP release as a rapid signal linking transient neuronal hypoactivity to persistent microglial engagement at inhibitory synaptic sites.

### P2Y12 stabilizes functional microglial bulbous endings

P2Y12 is a principal nucleotide receptor that mediates microglial process extension toward extracellular nucleotides after tissue injury or neuronal hyperactivity (*10–13, 23*). We found P2Y12 deficiency did not eliminate initial ATP-associated process-tip swelling, but impaired directed recruitment, localized microglial Ca²⁺ activity, and the conversion of transient swellings into stable BEs. Future studies will be needed to determine additional local cues that cooperate with purinergic signaling to initiate BE-like swellings. One candidate is UDP-P2Y6 signaling, because P2Y6 activation is known to elevate microglial Ca²⁺ (*17*) and UDP is able to attract microglial processes (*35*). P2Y12 downstream pathways may promote local membrane-potential regulation and cytoskeletal remodeling at the process tip, enabling ATP-associated swellings to be guided toward PV bouton-proximal sites and maintained as compartmentalized Ca²⁺-active structures. Thus, P2Y12 signaling may convert transient exploratory swellings into persistent BEs by coordinating both target-directed recruitment and local process-tip stabilization.

Recent studies have shown that microglia can selectively interact with, displace, or remove inhibitory synapses (*19, 36–38*). We previously showed that microglial shielding of inhibitory synapses is associated with increased neuronal activity during emergence from anesthesia (*18*). The present findings provide further insight into the upstream spatial cue and stabilization mechanism underlying this selective synaptic shielding. Astrocytic ATP provides a local cue near PV boutons, while P2Y12-dependent stabilization allows synaptic shielding to persist into early emergence. ATP-associated BEs formed during anesthesia may therefore continue to attenuate perisomatic inhibitory input during early emergence, thereby producing transient disinhibition and promoting the rebound increase in neuronal activity.

### Two-step control by neuromodulatory and spatial cues

Previous in vivo imaging studies showed that reduced neuronal activity or anesthesia increases microglial process surveillance, in part through a decrease in noradrenergic tone and microglial β2-adrenergic receptor signaling (*14, 15*). This neuromodulatory mechanism can place microglia in an interaction-permissive state, consistent with our previous finding that microglia preferentially contact perisomatic inhibitory synapses during anesthesia and promote neuronal activity during emergence (*18*). However, a reduction in norepinephrine may be insufficient to account for the spatial precision needed to direct individual microglial processes toward specific synaptic sites. Our current study supports a two-step model in which reduced noradrenergic signaling provides a permissive state cue that broadens microglial surveillance, whereas astrocyte-derived ATP provides a local positional cue that directs process recruitment and stabilizes P2Y12-dependent bulbous endings near perisomatic inhibitory inputs. The cooperation of these global neuromodulatory and local purinergic signals allows reduced circuit activity to be translated into synapse-selective microglial engagement, thereby facilitating rebound neuronal activity during emergence.

Recent studies have also implicated glial pathways in anesthetic-state regulation under distinct anesthetic regimens (*39, 40*), suggesting that glial responses are a broader component of anesthetic-state transitions. In our study, chemogenetic suppression of neuronal activity induced local ATP release and microglial BE formation, indicating that these responses are not solely attributable to a direct pharmacological effect of isoflurane. Increased ATP events and BE formation were also observed during natural sleep, consistent with previous reports of state-dependent changes in microglial surveillance, microglia–synapse interactions, and Ca²⁺ signaling during sleep (*29, 30, 41, 42*). Together, these observations suggest that local ATP release and microglial BE formation are associated with reduced circuit activity across experimentally induced and physiological brain states.

## Acknowledgments

This work is supported by the National Institutes of Health (R35NS132326 to L.-J.W. and R01AG041274 to Z.X.). Confocal microscopy was performed at the Center for Advanced Microscopy, a Nikon Center of Excellence, Department of Integrative Biology & Pharmacology at McGovern Medical School, UTHealth Houston (RRID: SCR_025962). We thank members of the Wu Lab and the Center for Neuroimmunology and Glial Biology for technical support and insightful discussions.

## Author contributions

K.H. and L.-J.W. conceived the project, designed the experiments, and wrote the manuscript. K.H., J.L., and S.M. performed animal surgery, in vivo image acquisition, behavioral experiments, immunofluorescence staining experiments, and data analyses. Y.L. and W.S. performed freely moving two-photon imaging experiments. D.K. performed acute slice imaging experiments. S.Z. assisted with P2Y12 cKO experiments. A.D.U. performed Cx3cr1-GCaMP7s-tdTomato mouse validation experiments. Y.U.L. and Z.X. provided expert advice and manuscript edits. M.J. contributed to experimental design, Panx1 conditional knockout-related experiments, and manuscript edits. L.-J.W. secured funding and supervised the project.

## Data availability

Source data are provided with the paper. The raw data that support the findings of this study are available from the corresponding author upon request.

## Code availability

All custom-written ImageJ macros and MATLAB scripts for analysis will be made available on request.

## Materials and Methods

### Animals

Male and female mice, 2 to 5 months of age, were used in accordance with institutional guidelines. All experimental procedures were approved by the Mayo Clinic Institutional Animal Care and Use Committee (IACUC, #5285-20, #7076-23), the Chinese Institute for Brain Research (#CIBR-IACUC 007) and the UTHealth Houston IACUC (#AWC-24-0046), depending on the location where the work was conducted, and were conducted in accordance with the NIH Guide for the Care and Use of Laboratory Animals. Mice were housed with a standard 12-h light (6:00 a.m. to 6:00 p.m.)/12-h dark (6:00 p.m. to 6:00 a.m.) at 22°C with 40-60% humidity and fed standard chow ad libitum. Neuron Ca²⁺ activity and locomotion activity were studied in C57BL/6 wild-type (WT) mice (JAX: 000664) using adeno-associated viruses (AAV). Cx3cr1^GFP/+^ (JAX: 005582) mice were used to visualize microglia-neuron interactions for in vivo two-photon imaging, with neurons labeled using AAV transfection. Cx3cr1^GFP/+^ mice were also crossed with PV-Cre knock-in mice (JAX:017320) for microglia and PV neuron interaction studies. Astrocytic Panx1 conditional knockout mice were achieved by crossing Panx1^fl/fl^ mice (GemPharmatech: T008074) with Aldh1l1^CreERT2^ mice (JAX: 031008). To visualize astrocytes and microglia simultaneously, Aldh1l1^CreERT2^ mice (JAX: 031008) were crossed with Ai14 (Rosa-CAG-LSL-tdTomato) reporter mice (JAX: 007914) and Cx3cr1^GFP/+^ mice to generate Aldh1l1^CreERT2^::Ai14::Cx3cr1^GFP/+^ mice. Cx3cr1-GCaMP7s-tdTomato mice, in which microglia are labeled with GCaMP7s and tdTomato, were kindly provided by Dr. Yong U. Liu at Guangzhou First People’s Hospital and were used to monitor microglial Ca²⁺ activity and morphology in vivo. To examine microglial Ca²⁺ activity in P2Y12-deficient mice, P2Y12 global knockout mice, kindly provided by Dr. Michael Dailey at the University of Iowa, were crossed with Cx3cr1-GCaMP7s-tdTomato mice. P2Y12^fl/fl^ mice were generated by Biocytogen Co., Ltd. and reported previously (Peng et al., Mol. Brain, 2019). Cx3cr1^CreER/+^ (JAX: 021160) mice were bred with P2Y12^fl/fl^ mice and Ai14 mice (JAX: 007914) to generate mice, in which P2Y12 was selectively deleted in microglia. Animals used for all experiments were randomly selected.

### Cranial window implantation and stereotaxic AAV injection

Under isoflurane anesthesia (3% induction, 1.5% maintenance), AAVs were injected into the motor cortex (anteroposterior: –1.0 mm, mediolateral: +1.0 mm) using a glass pipette and micropump (World Precision Instruments). AAVs were targeted to cortical layer II/III (dorsoventral: −0.2 to −0.3 mm from the cortical surface). A 300 nl volume was dispensed at a 50 nl/min rate, followed by a 5-min waiting period for diffusion. To image neuronal calcium activity, 300 nl of AAV9.CaMKII.GCaMP6s (#107790-AAV9, Addgene) or AAV2/9-CaMKIIa-jRGECO1a-WPRE-hGH-pA (#PT-3349, BrainVTA) was injected into the M1 cortex for in vivo two-photon imaging. To image ATP dynamics, AAV2/9-hSyn-ATP1.0 (#PT-1350, BrainVTA), encoding GRAB-ATP1.0, was injected into the motor cortex. To study the relationship between microglia-neuron interactions and neuron Ca²⁺ activity, we injected a cocktail of three viruses: AAV1.CaMKII.Cre (#105558-AAV1, Addgene), AAV9.Syn.Flex.GCaMP6s (#100845-AAV9, Addgene), and AAV9.CAG.FLEX.tdTomato (#28306-AAV9, Addgene), or AAV2/9.CaMKII.jRGECO1a (#PT-3349, BrainVTA) into M1 cortex of Cx3cr1^GFP/+^ mice as indicated for each experiment. To visualize microglia and PV boutons simultaneously, we injected AAV9.CAG.FLEX.tdTomato into M1 cortex of Cx3cr1^GFP/+^::PV^Cre/+^ mice. To visualize astrocytic Ca²⁺ activity and PV boutons, we injected AAV2/9-GfaABC1d-GCaMP6f (#PT-2560, BrainVTA) and AAV9.CAG.FLEX.tdTomato into PV^Cre/+^ mice. To manipulate CaMKII neuron activity, we injected AAV1.CaMKII.Cre (#105558-AAV1, Addgene) and AAV2-hSyn-DIO-hM4D(Gi)-mCherry (#44362-AAV2, Addgene) into Cx3cr1^GFP/+^ mice.

Mice were implanted with a cranial window immediately after virus injection. A 3 mm circular coverslip was sealed in place with light-curing dental cement (Tetric EvoFlow, Ivoclar). The skull, excluding the region with the window, was then covered with iBond Total Etch primer (Heraeus) and cured with LED light. Finally, a head bar (Neurotar) was attached to the skull using light-curing dental glue, and all exposed skull surfaces were also covered by dental glue. Mice were allowed to recover from anesthesia on a heating pad for 10 min before they were returned to their home cage. Ibuprofen was provided in drinking water as an analgesic for 72 h before and after surgery.

### Drug administration

The effect of Panx1 inhibition was tested using probenecid (PBN; #P36400, Invitrogen) at 200 mg/kg body weight. PBN was dissolved in 0.9% sterile saline and injected intraperitoneally 4 h before two-photon imaging. For chemogenetic neuronal silencing experiments, deschloroclozapine (DCZ; #HY-42110, MedChemExpress) was administered intraperitoneally at 100 μg/kg during in vivo imaging. To induce CreER-mediated recombination, tamoxifen (#T5648, MilliporeSigma) was administered intraperitoneally at 150 mg/kg after dissolution in corn oil at 20 mg/ml. For P2Y12 cKO mice, tamoxifen administration was initiated 6 weeks before imaging and 3 weeks before cranial window surgery, with eight injections in total spaced 48 h apart. For Panx1 cKO mice, tamoxifen administration was initiated 30 days before imaging or tissue collection and was given once daily for six consecutive days.

### In vivo two-photon imaging

After recovery from virus injection and cranial window surgery (2–4 weeks), mice were trained to move on an air-lifted platform (NTR000251-06, Neurotar) while head-fixed under a two-photon microscope for 30 min/day during the 3 days before imaging. Across all studies, mice were allowed 10 min to acclimate after being placed in the head restraint before imaging began. Mice were imaged using a two-photon imaging system (Scientifica) equipped with a tunable laser (Chameleon Vision II; Coherent). Laser wavelength was tuned to 940 nm to image Cx3cr1^GFP/+^ microglia or ATP sensor fluorescence, GCaMP6s fluorescence and tdTomato-labelled neurons or microglia simultaneously. Imaging utilized a 20× water-immersion lens and a 320×320 μm field of view (1024 × 1024 pixels). The microscope was equipped with a 565 nm dichroic mirror and the following emission filters: 525/50 nm (green channel) and 620/60 nm (red channel) for GFP/tdTomato imaging. The laser power was maintained at 30–40 mW under the objective. Imaging in the cortex was conducted at a depth of 100–200 μm (Layer II/III).

To visualize cortical blood vessels during in vivo imaging, Texas Red-conjugated 70-kDa dextran was injected intravenously through the tail vein shortly before imaging (50 μl, 20 mg/ml; #D1864, Thermo Fisher Scientific).

For chemogenetic neuronal silencing experiments, Cx3cr1^GFP/+^ mice expressing GRAB-ATP1.0 together with hM4D(Gi)-mCherry or tdTomato in cortical CaMKII neurons were imaged under awake head-fixed conditions 3–4 weeks after AAV injection. A 120-min baseline period was recorded before DCZ administration, followed by continuous imaging for 4 h at 1 frame/s. ATP sensor fluorescence and microglial GFP were acquired in the green channel, and hM4D(Gi)-mCherry or tdTomato was acquired in the red channel.

To image neuronal Ca²⁺ activity, we acquired T-series (1.0 Hz frame rates, 512 × 512 pixels). Mice were first imaged under awake baseline conditions (15 min). A nose cone was then secured against the head-restraint system and the mouse was induced with isoflurane on the platform (3%). Isoflurane was maintained at 1.0–1.5% during the 30-min anesthesia imaging period. Under isoflurane, body temperature was maintained at 37°C using a heating pad. To end anesthesia, the nose cone was removed, and mice were imaged during this recovery period (‘emergence from anesthesia’) over four 15 min blocks (60 min emergence). Mouse locomotion was recorded during in vivo imaging using the Mobile HomeCage magnetic tracking system (NTR000251-06 and NTR000293-02, Neurotar).

To image ATP dynamics or microglial Ca²⁺ activity, we acquired T-series (0.5-1.0 Hz frame rates, 1024 × 1024 pixels). To image microglia-neuron interactions, we also acquired Z-stacks (11 sections, 2 μm step size, 1024 × 1024 pixels) once every 1-2 min. Imaging data were excluded from analysis only in the instance of technical failure. For ATP imaging in astrocytic Panx1 cKO mice, in vivo two-photon imaging was performed 2–4 weeks after AAV injection using an Olympus Fluoview FV4000-MPE microscope coupled to a Newport Insight Deep See Ti laser. GRAB-ATP1.0 fluorescence was imaged in layer II/III of the motor cortex at a depth of 150–200 μm using a 25× water-immersion objective (XLPLN25XWMP2, NA 1.05; Olympus). Images were acquired at 1.5 Hz with a field of view of 320 × 320 μm and a resolution of 1024 × 1024 pixels.

### Acute cortical slice imaging

Acute cortical slices were prepared from Cx3cr1^GFP/+^ mice to examine microglial process responses to local ATP or GABA application. Mice were deeply anesthetized and decapitated. Brains were rapidly removed and placed in ice-cold oxygenated artificial cerebrospinal fluid (ACSF) bubbled with 95% O_2_ and 5% CO_2_. ACSF contained the following components in mM: 126 NaCl, 26 NaHCO_3_, 2.5 KCl, 1.25 NaH_2_PO_4_, 2 CaCl_2_, 2 MgCl_2_, and 10 glucose, with sucrose added to adjust osmolarity to 300–320 mOsm. Coronal cortical slices were cut at 300 μm thickness in ice-cold ACSF using a vibrating blade microtome (VT1000S, Leica) and transferred to a recovery chamber containing oxygenated ACSF at room temperature for at least 30 min before imaging.

Slice imaging was performed at room temperature in a perfusion chamber continuously supplied with oxygenated ACSF at approximately 2 ml/min. Microglia were imaged 50–120 μm below the slice surface using the same two-photon microscope and detection configuration described above.

For local pharmacological application, a glass pipette was filled with ATP (1 mM) or GABA (100 μM or 1 mM) together with Alexa Fluor 594 dissolved in ACSF to visualize the pipette tip and local diffusion. The pipette tip was positioned under two-photon guidance near cortical microglial processes, with care taken to avoid direct contact with microglial somata or major processes. For ATP application, microglial process dynamics were monitored after pipette insertion, allowing ATP to diffuse passively from the pipette tip. For GABA application, microglial process dynamics were first monitored for 20 min after pipette insertion during passive diffusion, followed by pressure-puff stimulation through the same pipette using a Precision Pneumatic Picopump Microinjector (PV820, WPI).

Time-lapse images were acquired at 0.5–1 Hz with a resolution of 1024 × 1024 pixels to monitor microglial process dynamics around the pipette tip. Process accumulation was quantified as GFP mean fluorescence intensity (MFI) within a 50-μm-radius ROI centered on the pipette tip, and processes inside and outside this ROI were defined as proximal and distal processes, respectively.

### Imaging data analysis

#### Image processing

Z-stack and time-series images were corrected for lateral motion and focal-plane displacement using ImageJ (National Institutes of Health, version 1.53f51) with the StackReg and TurboReg plugins (*43*). For representative images, z-stack images were displayed as maximum- or average-intensity projections, and time-series images were displayed as average-intensity projections over the indicated time window.

#### Neuronal calcium signal processing and quantification

Neuronal Ca²⁺ imaging data were analyzed using ImageJ and custom MATLAB scripts. For ROI selection, an average-intensity image was generated from each time-series video, and ROIs were manually drawn around neuronal somata in cortical layer II/III using the oval selection tool in ImageJ. Neurons that did not show clear somatic GCaMP6s fluorescence, characterized by a bright cytoplasmic ring and dark nucleus during the awake baseline period, were excluded from analysis. MFI traces were extracted from each ROI and analyzed within defined imaging windows corresponding to awake baseline, isoflurane anesthesia, and emergence from anesthesia.

For each ROI, fluorescence traces were smoothed using a Gaussian filter with a 3-frame window. Baseline fluorescence (F0) was defined as the 25th percentile of the smoothed fluorescence trace within each analysis window, and ΔF/F was calculated as (F − F0)/F0. ROIs with non-finite or zero baseline values were excluded from analysis. The maximum amplitude was defined as the maximum ΔF/F value within each analysis window. Time active was calculated as the cumulative time during which ΔF/F exceeded 0.25. Signal area was calculated as the integrated ΔF/F signal above this threshold.

For state comparisons, Ca²⁺ activity parameters were averaged across ROIs within each mouse and then compared across experimental conditions. Neurons were classified as showing increased activity when signal area was greater than 1.5-fold the awake baseline value and decreased activity when signal area was less than 0.5-fold the awake baseline value. For analyses comparing neuronal Ca²⁺ activity with locomotor activity, locomotion speed was synchronized to imaging frames and summarized within the same analysis windows used for Ca²⁺ imaging. The relationship between neuronal Ca²⁺ activity and locomotion speed was assessed using simple linear regression.

#### GRAB-ATP event detection and quantification

GRAB-ATP1.0 signals were analyzed using AQuA 2.0 (*44*) or custom ImageJ scripts, depending on the experimental design. For automated event detection, pixel-wise ΔF/F images were generated, and local fluorescence maxima exceeding a ΔF/F threshold of 0.2 were identified as ATP release hotspots. Each detected hotspot was fitted with a two-dimensional Gaussian to determine its centroid and area. Events detected within a 10-frame temporal window and a 5-μm spatial radius were merged and counted as a single event. ATP hotspot frequency, peak amplitude, area, and spatial coordinates were extracted for subsequent analyses.

For experiments requiring direct alignment of ATP signals with microglial morphology, including simultaneous imaging of GRAB-ATP1.0 and Cx3cr1-GFP microglia, regions of interest were manually placed over transient ATP signals in ImageJ, and ΔF/F traces were extracted to quantify ATP dynamics relative to microglial process movement and bulbous-ending formation.

#### Detection of microglial bulbous endings

Bulbous endings (BEs) were defined as spherical enlargements at the terminal endings of Cx3cr1-GFP microglial processes that persisted for ≥5 min and showed a diameter greater than 2-fold of the adjacent process shaft. BE-like swellings were defined using the same morphological criteria but persisted for <5 min. Image stacks were inspected at 1-min intervals, and each BE or BE-like swelling was confirmed by morphological continuity with the originating microglial process. For ATP-associated BE analysis, a stable BE was classified as ATP-associated when it formed within an ATP hotspot ROI within 10 min after ATP event onset and persisted for ≥5 min.

#### Microglial contact and process motility analysis

Microglia-neuron contacts were analyzed from time-series images, with z-stack images used when available to confirm BE morphology and process continuity. For analyses restricted to soma- or PV bouton-contacting BEs, contact was defined when the closest distance between the Cx3cr1-GFP signal at the BE and the neuronal soma or PV bouton fluorescence signal was less than 1 μm in the same focal plane.

Microglial process motility was quantified from maximum-intensity projections of time-lapse images by manually tracking primary process tips in ImageJ using the Manual Tracking plugin. Process directness was calculated as the Euclidean displacement of each process tip divided by the accumulated path length.

#### Spatial proximity and nearest-neighbor analysis

To determine whether spatial relationships among PV boutons and ATP spots, Panx1 puncta and synaptic markers, and astrocytic Ca²⁺ events and PV boutons exceeded chance levels, we performed randomization-based nearest-neighbor analysis. For each dataset, centroid coordinates were obtained from manually defined ROIs or from puncta detected after thresholding using the Particle Analyzer function in ImageJ; astrocytic Ca²⁺ event centroids were obtained from AQuA2.0. Nearest-neighbor distances were calculated after coordinate alignment and conversion to physical units in micrometers. For ATP spot-to-PV bouton and astrocytic Ca²⁺ event-to-PV bouton analyses, experimentally observed PV bouton positions were kept fixed while ATP spot or Ca²⁺ event positions were randomly redistributed within the same field of view. For Panx1-to-synaptic marker analyses, experimentally observed synaptic marker positions were kept fixed while Panx1 puncta were randomly redistributed within the same field of view. Randomization was repeated for 1,000 iterations, and nearest-neighbor distances were recalculated for each iteration and compared with the observed nearest-neighbor distances. Summary measures, including mean nearest-neighbor distance, median nearest-neighbor distance, and the fraction of points within an analysis-specific distance threshold, were calculated for each dataset and used for group-level statistical analyses. To assess whether stable BE formation was spatially enriched at ATP hotspots, observed ATP hotspot ROIs were compared with matched randomized control ROIs. For each observed hotspot, five circular control ROIs of identical size were placed at nonoverlapping locations within the same field of view and matched approximately for distance to the nearest PV bouton, local microglial process density, and assigned event onset time. Control ROIs did not overlap with detected ATP hotspots. The percentage of observed and randomized ROIs associated with stable BE formation was calculated for each mouse.

### Astrocytic/microglial Ca²⁺ event analysis

Astrocytic and microglial Ca²⁺ events were detected from motion-corrected time-lapse images using AQuA2.0 implemented in MATLAB. For image datasets, the central 950×950-pixel region of each 1024 × 1024-pixel image was cropped and downsampled to 475×475 pixels before AQuA2.0 analysis. Temporal and spatial sampling parameters were set according to the acquisition settings for each dataset. Significant fluorescence transients were automatically detected using the following parameters: intensity threshold scaling factor, 3.0; minimum duration, 5 frames; minimum area, 50 pixels; seed significance Z-score, 3.5; and source-level sensitivity, 8. Spatial segmentation was enabled, whereas decay speed calculation and network feature analysis were disabled. Automatically detected Ca²⁺ event ROIs were manually validated before quantification. For each detected event, the centroid, spatial area, circularity, duration, peak amplitude, and timing were extracted. Event frequency was calculated for each imaging period. Spatial area was defined as the two-dimensional area occupied by each detected Ca²⁺ event mask. Astrocytic Ca²⁺ event centroids were used for nearest-neighbor analysis with PV boutons after coordinate alignment and conversion to physical units. Microglial Ca²⁺ events were assigned to bulbous endings or adjacent processes based on spatial overlap with tdTomato-labeled microglial morphology.

#### Immunohistochemistry

Mice were deeply anesthetized with isoflurane and transcardially perfused with phosphate-buffered saline (PBS) for 1 min, followed by 4% paraformaldehyde (PFA). Brains were post-fixed overnight in 4% PFA and immersed in 30% sucrose for 2 days for cryoprotection. The brains were sectioned into 30 μm slices. After PBS wash, slices were incubated for 1 hour in 5% bovine serum albumin (BSA) and 0.5% Triton X-100 dissolved in PBS, followed by incubation overnight at 4°C with primary antibodies diluted in blocking solution: anti-IBA1 (1:500, #ab5076, Abcam), anti-GFAP (1:500, #3670S, Cell Signaling Technology), anti-VGAT (1:500, #131 004, Synaptic Systems), anti-VGLUT1 (1:500, #135 304, Synaptic Systems), anti-Panx1 (1:200, #488100, Thermo Fisher Scientific), anti-NeuN (1:500, #ab104225, Abcam), anti-P2Y12 (1:500, #848002, BioLegend). Slices were then incubated for 3 h at room temperature with secondary antibodies diluted in blocking solution: anti-goat Alexa Fluor 488, anti-rabbit Alexa Fluor 647, anti-rabbit Alexa Fluor 405, anti-mouse Alexa Fluor 647, anti-rat Alexa Fluor 650 (1:500, #A32814, #A32795, #A48258, #A32787, # SA5-10029, Thermo Fisher Scientific), and anti-guinea pig Cy3 (1:500, 706-165-148, Jackson ImmunoResearch Laboratories). Slices were mounted on glass slides with Fluoromount-G (00-4958-02, Invitrogen).

For Panx1 expression analysis in astrocytes, 50-μm motor cortex sections were permeabilized with 0.5% Triton X-100, blocked for 1 h in PBS containing 5% BSA and 0.3% Triton X-100, and incubated overnight at 4°C with the anti-Panx1 antibody described above and anti-GFAP antibody (1:1,000, #ab4674, Abcam). Sections were then incubated for 2 h at room temperature with fluorophore-conjugated secondary antibodies and mounted with DAPI (0.1 μg/ml) in 50% glycerol/PBS.

Confocal images were acquired using a Nikon AXR microscope with NIS-Elements software version 6.20. Panx1 expression images were acquired using a Leica TCS SP8 confocal microscope and analyzed with Imaris (v10.1, Oxford Instruments) or ImageJ. For P2Y12 distribution analysis, images were acquired using a Nikon CSU-W1 SoRa super-resolution spinning-disk system with a 100× objective and analyzed using the segmented line tool in ImageJ.

#### EEG–EMG electrode implantation

For experiments combining freely moving miniature two-photon imaging with EEG–EMG recordings, EEG and EMG electrodes were implanted during the cranial window surgery. After placement of the cranial window, stainless steel screw electrodes were inserted into the frontal and parietal skull regions for EEG recording, and additional screws were used for grounding. Insulated silver wires were inserted into the neck muscles for EMG recording. The EEG–EMG electrode assembly and miniature microscope baseplate were secured to the skull using light-curing dental cement. Mice were allowed to recover for at least one week before recording and were habituated to the recording cage and cable connection before imaging.

#### Freely moving miniature two-photon imaging

For freely moving miniature two-photon imaging, mice expressing GRAB-ATP1.0 in the cortex were used after recovery from cranial window implantation and EEG–EMG electrode placement. Before imaging, mice were briefly head-fixed to identify the imaging field and to attach the miniature microscope probe to the implanted baseplate. Mice were then placed in the recording cage, and EEG–EMG electrodes were connected to the recording system.

Imaging was performed using a miniature two-photon imaging system (SUPERNOVA-100, headpiece FHIRM-U; TransVista) controlled by SUPERGIN imaging software. GRAB-ATP fluorescence and Cx3cr1^GFP/+^ microglia were imaged using a 920-nm femtosecond fiber laser. Images were acquired at 5 Hz with a frame size of 1024 × 880 pixels. The average laser power delivered to the brain was maintained below 60 mW. Imaging sessions were performed during the light and dark phases to monitor ATP dynamics and microglial morphology across natural sleep–wake states.

Imaging periods were aligned to EEG–EMG-defined behavioral states. For light- and dark-phase analyses, ATP hotspots were quantified from imaging periods recorded during the corresponding phase. For vigilance-state-specific analyses, wake and NREM sleep episodes were selected based on EEG–EMG classification. ATP hotspots were detected using the same GRAB-ATP analysis pipeline used for head-fixed in vivo imaging. ATP hotspot frequency and spot area were quantified for each imaging session. Microglial BEs were identified from Cx3cr1^GFP/+^ time-lapse images using the same morphological criteria used for head-fixed imaging, and BE number was quantified during wake and NREM sleep episodes.

#### Sleep-state classification and sleep imaging analysis

EEG–EMG recording was initiated before each miniature two-photon imaging session, and the start time of imaging was marked in the EEG–EMG recording file for temporal alignment. Cortical EEG and neck EMG signals were acquired at 250 Hz using the Medusa system (Bio-Signal Technologies). Polygraphic recordings were segmented into wakefulness, NREM sleep, or REM sleep in 4-s or 10-s epochs according to standard EEG–EMG criteria. Delta (0.5–4 Hz), theta (4–10 Hz), and sigma (10–15 Hz) band powers were calculated from the EEG signal. The EMG envelope was obtained by rectification followed by low-pass filtering. Wakefulness was defined by high EMG activity with low delta power, NREM sleep by high delta power with low-to-moderate EMG activity, and REM sleep by low EMG activity with an elevated theta/delta power ratio. Automatically assigned sleep–wake stages were visually inspected and manually corrected when necessary.

#### Data analysis and statistics

The experimental unit was defined according to the experimental design. Individual animals were treated as independent biological replicates for mouse-level comparisons. For analyses involving individual cells, ATP events, bulbous endings, regions of interest, or synaptic puncta, the unit of analysis and sample size are indicated in the corresponding figure legends. No statistical methods were used to predetermine sample sizes, but sample sizes were chosen based on previous studies using similar in vivo imaging and histological analyses (*14, 18*). Animals were randomly assigned to experimental groups when applicable. Wherever possible, image quantification or event classification was confirmed by a second investigator blinded to experimental condition or genotype. Data were excluded only in cases of technical failure, such as poor viral expression, loss of the cranial window, or failure to recover the same imaging field.

Data were analyzed using GraphPad Prism 10. The experimental unit and sample size for each analysis are indicated in the corresponding figure legends. For analyses involving multiple cells, ATP events, bulbous endings, regions of interest, or synaptic puncta from the same animal, the number of observations and the number of animals are reported where applicable. Comparisons between two groups were performed using two-sided paired or unpaired t-tests, as appropriate. Comparisons among multiple groups were performed using one-way ANOVA, repeated-measures ANOVA, two-way ANOVA, or mixed-effects analysis followed by Tukey’s, Dunnett’s, or Fisher’s LSD multiple-comparisons test, as indicated in the figure legends. Categorical data were analyzed using Fisher’s exact test. Relationships between two continuous variables were analyzed using simple linear regression. Data are presented as mean ± SEM. Differences were considered statistically significant at P < 0.05.

---

**Supplementary Fig. 1.**
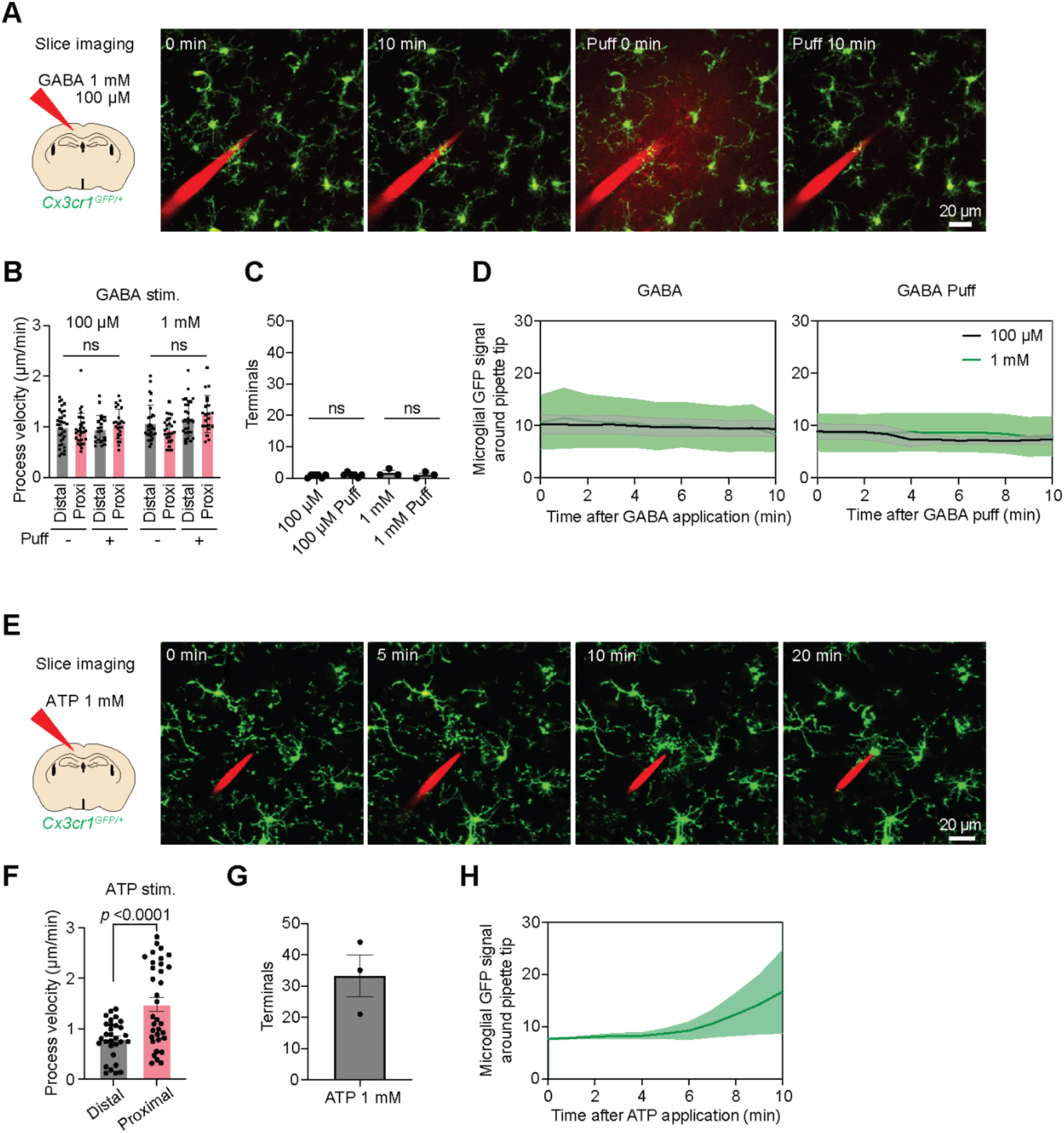
ATP, but not GABA, rapidly recruits microglial processes in acute cortical slices. **(A)** Representative two-photon images of Cx3cr1^GFP/+^ microglia in an acute cortical slice during local GABA diffusion and subsequent puff stimulation. GABA was applied through a glass pipette. **(B)** Quantification of microglial process tip velocity in distal and proximal processes before and after GABA puff application (two-way ANOVA with Tukey’s multiple-comparisons test; n = 21–34 processes per group from 3 mice). **(C)** Quantification of the number of microglial process tips recruited to the GABA pipette before and after puff stimulation (two-way ANOVA with Tukey’s multiple-comparisons test; n = 3 to 6 slices per group from 3 mice). **(D)** Time courses of microglial GFP signal around the GABA pipette after local GABA application or puff stimulation. **(E)** Representative two-photon images of Cx3cr1^GFP/+^ microglia in an acute cortical slice during local ATP application. ATP was applied through a glass pipette. **(F)** Quantification of microglial process tip velocity after ATP stimulation. Process tips proximal to the ATP pipette were compared with distal process tips (two-sided unpaired t-test; distal, n = 30 processes; proximal, n = 34 processes; data collected from 3 mice). **(G)** Quantification of the number of microglial process tips recruited to the ATP pipette. **(H)** Time course of microglial GFP signal around the ATP pipette after ATP application. Data are mean ± SEM. Individual points represent process tips or slices as indicated. ns, not significant.

**Supplementary Fig. 2.**
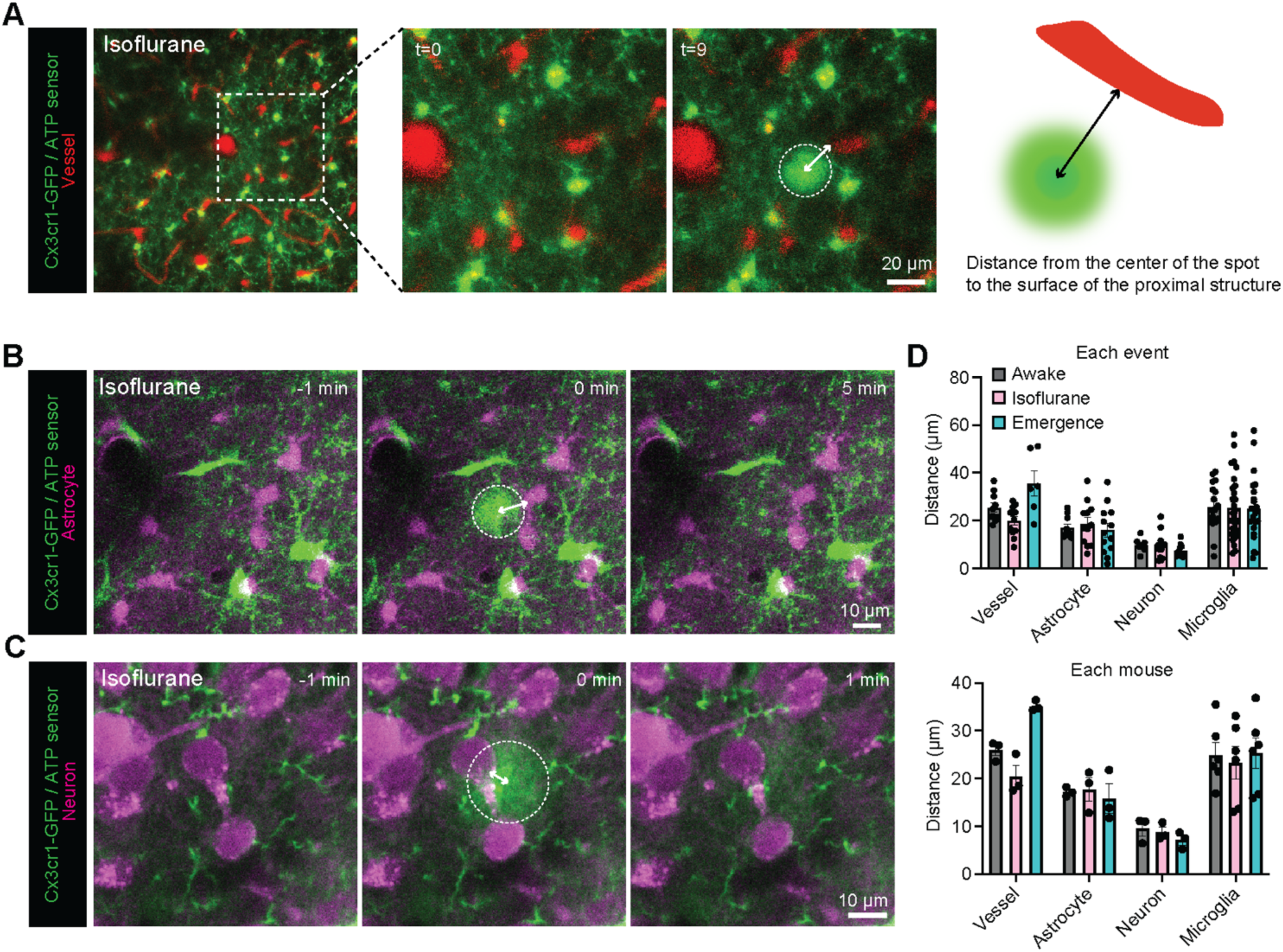
Spatial relationship between ATP hotspots and cortical structures during anesthesia. **(A)** Representative in vivo two-photon images showing ATP hotspots, microglia, and cortical blood vessels during isoflurane anesthesia. ATP was monitored with GRAB-ATP1.0 in Cx3cr1^GFP/+^ mice, and blood vessels were visualized with Texas Red-conjugated 70-kDa dextran. Dashed circles indicate ATP hotspots. The schematic illustrates distance measurement from the center of an ATP hotspot to the surface of the nearest cortical structure. **(B)** Representative in vivo images showing ATP hotspots, microglia, and astrocytes during isoflurane anesthesia. Astrocytes were genetically labeled with tdTomato using Aldh1l1^CreERT2^::Ai14 (Rosa-CAG-LSL-tdTomato) mice crossed with Cx3cr1^GFP/+^ mice. **(C)** Representative in vivo images showing ATP hotspots, microglia, and excitatory neuronal somata during isoflurane anesthesia. Excitatory neurons were labeled with tdTomato by co-injection of AAV-CaMKII-Cre and AAV-CAG-FLEX-tdTomato into the motor cortex of Cx3cr1^GFP/+^ mice. **(D)** Quantification of the distance from ATP hotspots to blood vessels, astrocytes, neuronal somata, and microglia across awake baseline, isoflurane anesthesia, and emergence. Distances are shown for individual ATP events and as mouse averages. Data are shown as mean ± SEM with individual ATP events or mice shown as points.

**Supplementary Fig. 3.**
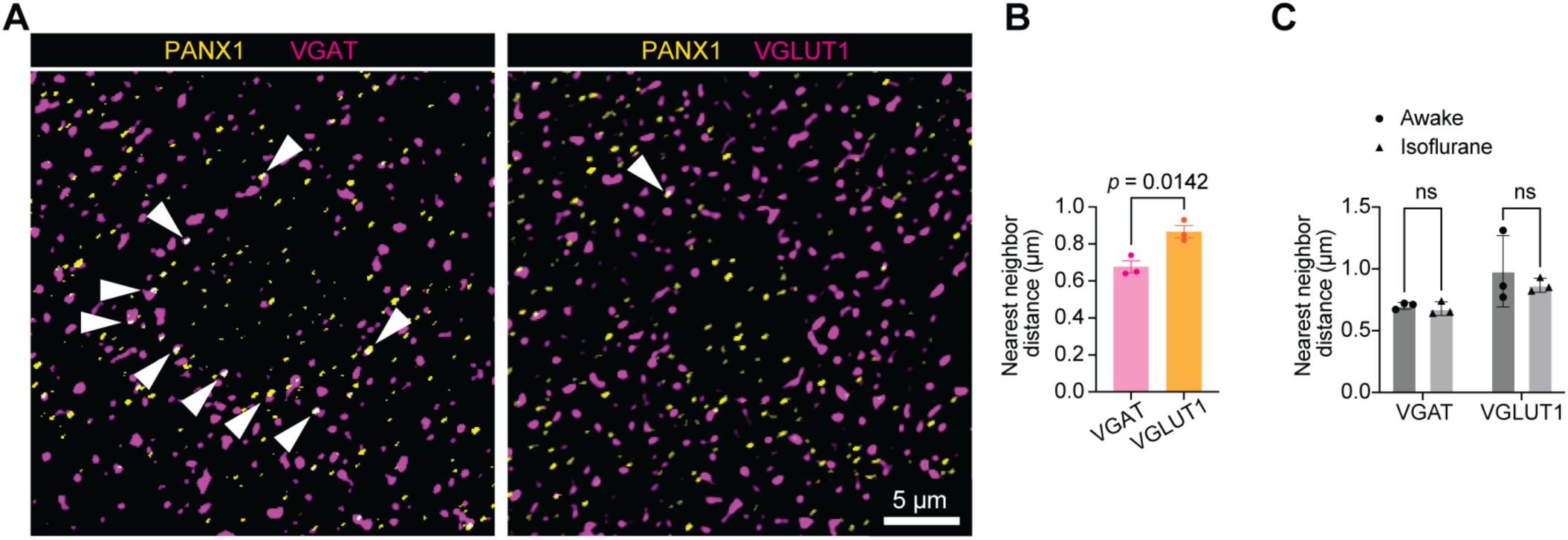
Spatial relationship between PANX1 and inhibitory or excitatory synaptic markers. **(A)** Representative confocal images showing PANX1 immunoreactivity together with VGAT-positive inhibitory synaptic markers or VGLUT1-positive excitatory synaptic markers in cortical tissue. Arrowheads indicate PANX1 puncta located near VGAT or VGLUT1 signals. **(B)** Comparison of observed nearest-neighbor distances between PANX1 and VGAT puncta or between PANX1 and VGLUT1 puncta (two-sided unpaired t-test; n = 3 mice). **(C)** Quantification of nearest-neighbor distances between PANX1 and VGAT or VGLUT1 puncta during awake and isoflurane anesthesia (two-way ANOVA with Fisher’s LSD test; n = 3 mice per group). Data are shown as mean ± SEM.

**Supplementary Fig. 4.**
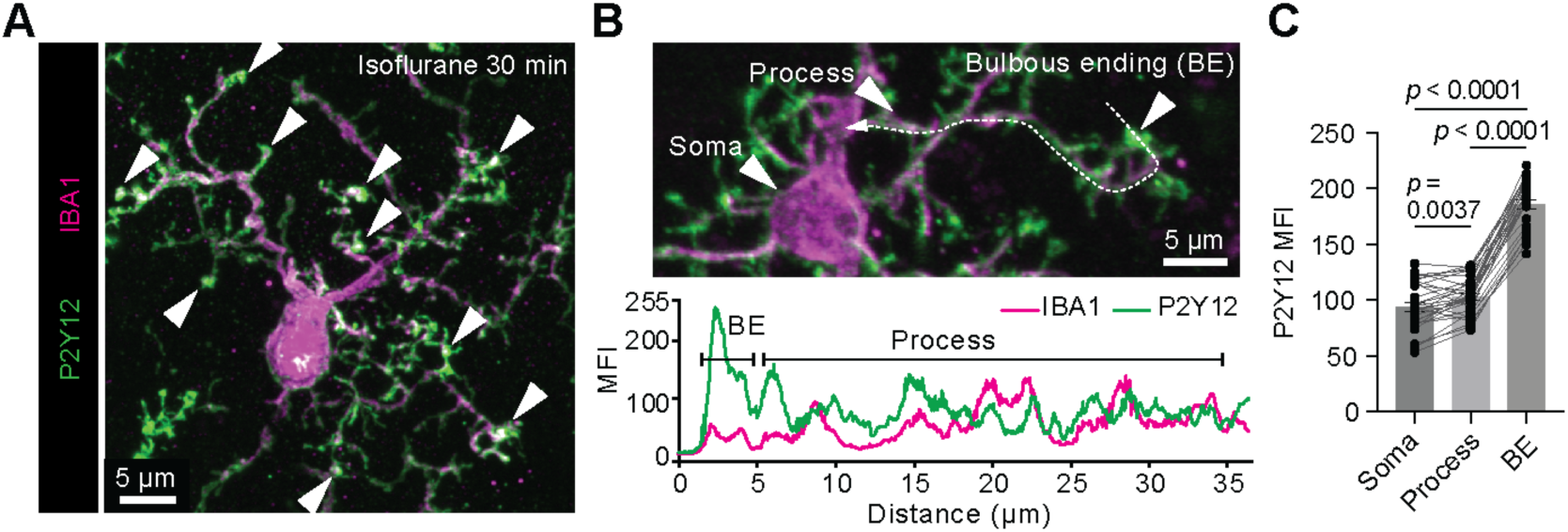
P2Y12 enrichment in microglial bulbous endings. **(A)** Representative super-resolution image showing IBA1-positive microglia and P2Y12 immunoreactivity in cortical tissue collected during isoflurane anesthesia. Arrowheads indicate bulbous endings (BEs). **(B)** Representative high-magnification image and corresponding fluorescence intensity profile of IBA1 and P2Y12 signals along a microglial soma–process–BE axis. The dashed line indicates the line ROI used for fluorescence intensity measurement. **(C)** Quantification of P2Y12 mean fluorescence intensity (MFI) in microglial somata, processes, and BEs. Individual lines connect measurements obtained from the same microglial cell (one-way repeated-measures ANOVA with Tukey’s multiple-comparisons test, n = 30 microglia from 3 mice). Data are shown as mean ± SEM. Individual points represent microglia. BE, bulbous ending.

**Supplementary Fig. 5.**
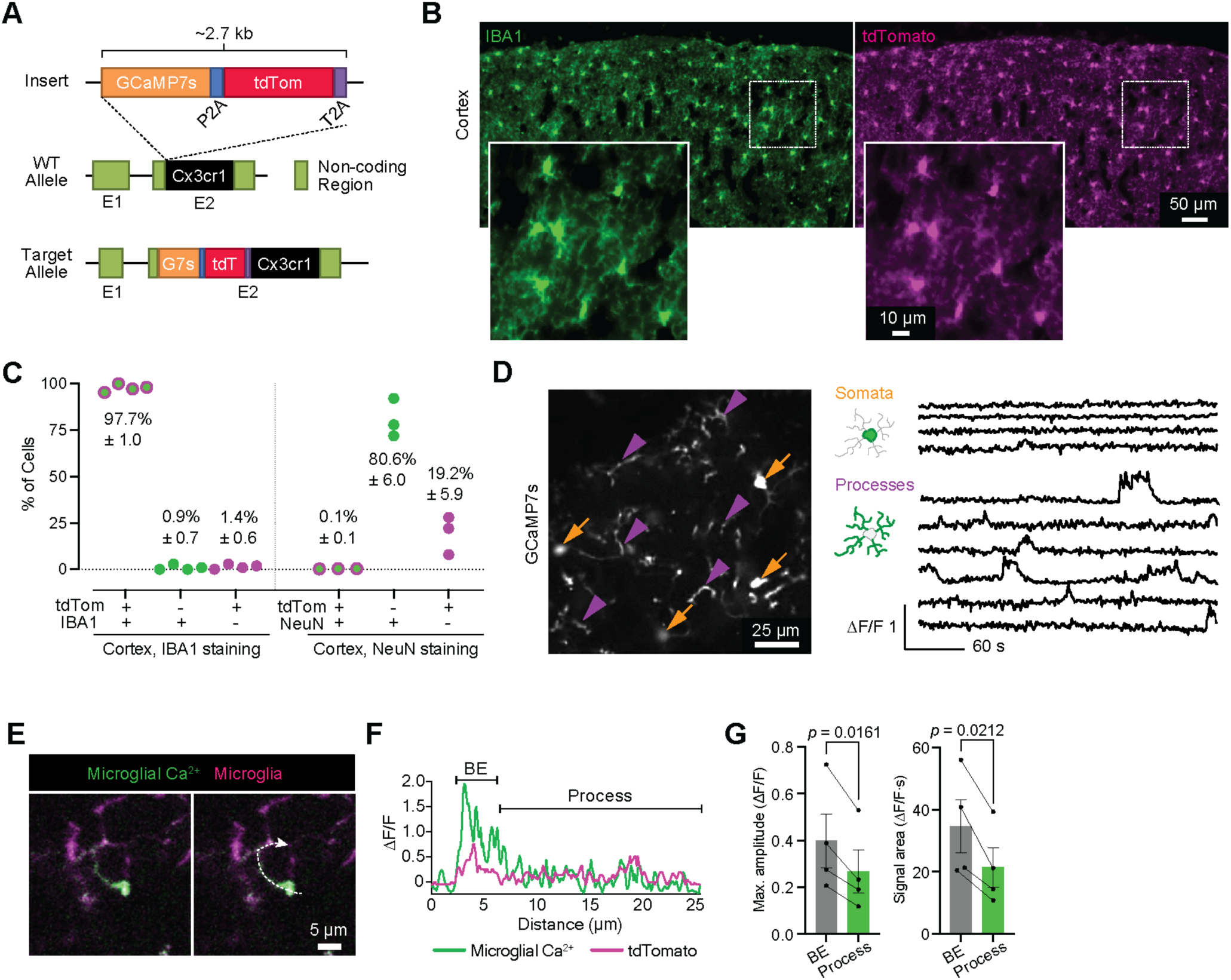
Characterization of the Cx3cr1-GCaMP7s-tdTomato mouse line and compartmentalized microglial Ca²⁺ activity in bulbous endings. (A) Gene-targeting strategy used to generate the Cx3cr1-GCaMP7s-tdTomato mouse line. The GCaMP7s-P2A-tdTomato-T2A cassette was inserted immediately upstream of the Cx3cr1 coding sequence, allowing constitutive expression of both GCaMP7s and tdTomato in microglia. (B) Representative cortical images showing endogenous tdTomato fluorescence together with IBA1 immunostaining to assess microglial labeling specificity. Insets show high-magnification views of the boxed regions. (C) Quantification of tdTomato labeling specificity in cortical tissue. Cells were classified according to tdTomato and IBA1 or NeuN immunoreactivity. The left graph shows the percentages of tdTomato⁺/IBA1⁺, tdTomato⁺/IBA1⁻, and tdTomato⁻/IBA1⁺ cells. The right graph shows the percentages of tdTomato⁺/NeuN⁺, tdTomato⁻/NeuN⁺, and tdTomato⁺/NeuN⁻ cells. IBA1 staining, n = 4 mice; NeuN staining, n = 3 mice. (D) Representative in vivo two-photon image and ΔF/F Ca²⁺ traces from microglial somata and processes reported by GCaMP7s in an awake mouse. Orange arrows indicate somatic Ca²⁺ events, and purple arrowheads indicate process Ca²⁺ events. (E) Representative in vivo two-photon images showing localized microglial Ca²⁺ activity within a BE. Microglial Ca²⁺ activity is shown in green, and microglial morphology is shown in magenta. Arrowheads indicate the BE. (F) Fluorescence intensity profiles of microglial GCaMP7s and tdTomato signals across a BE and the adjacent process. (G) Mouse-level quantification of microglial Ca²⁺ activity in BEs and adjacent processes, corresponding to the event-level analysis in Fig. 4F. Maximum amplitude and signal area are shown (two-sided paired t-test, n = 4 mice). Data are shown as mean ± SEM. Individual points represent mice. BE, bulbous ending.

**Supplementary Fig. 6.**
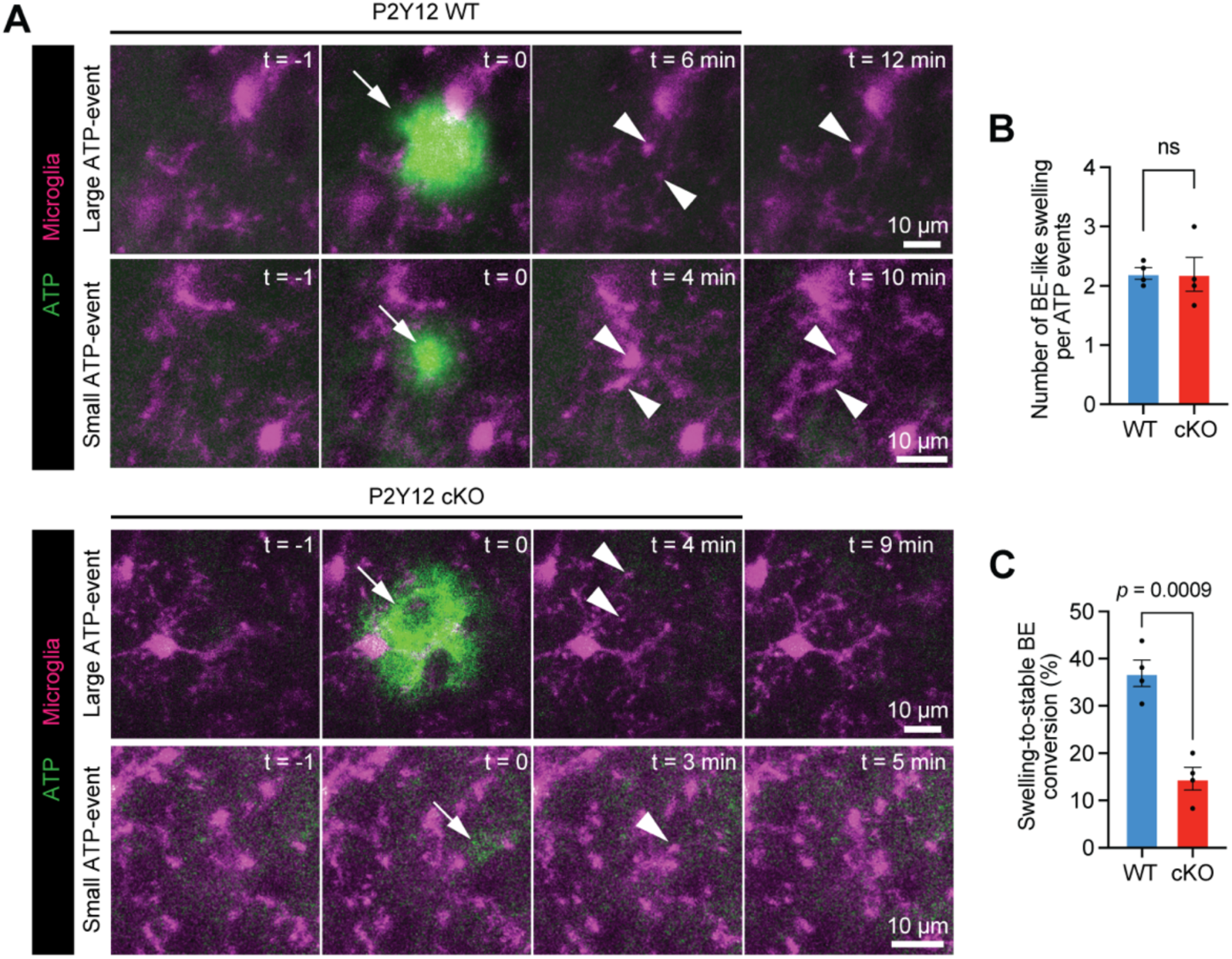
P2Y12 promotes stabilization of ATP-associated microglial bulbous endings. **(A)** Representative in vivo two-photon time-lapse images showing ATP-associated BE-like swellings in P2Y12 WT and microglia-specific P2Y12 cKO mice during isoflurane anesthesia. Large and small ATP events are shown separately. ATP signals are shown in green and microglial morphology in magenta. Arrows indicate ATP events, and arrowheads indicate BE-like swellings or stable BEs. **(B)** Quantification of the number of initial BE-like swellings formed per ATP event in P2Y12 WT and cKO mice (two-sided unpaired t-test, n = 4 mice per group). **(C)** Percentage of ATP-associated process-tip swellings that developed into stable BEs, defined as those persisting for ≥5 min, in P2Y12 WT and cKO mice (two-sided unpaired t-test, n = 4 mice per group). Data are shown as mean ± SEM. Individual points represent mice. BE, bulbous ending; cKO, conditional knockout; ns, not significant.

**Supplementary Fig. 7.**
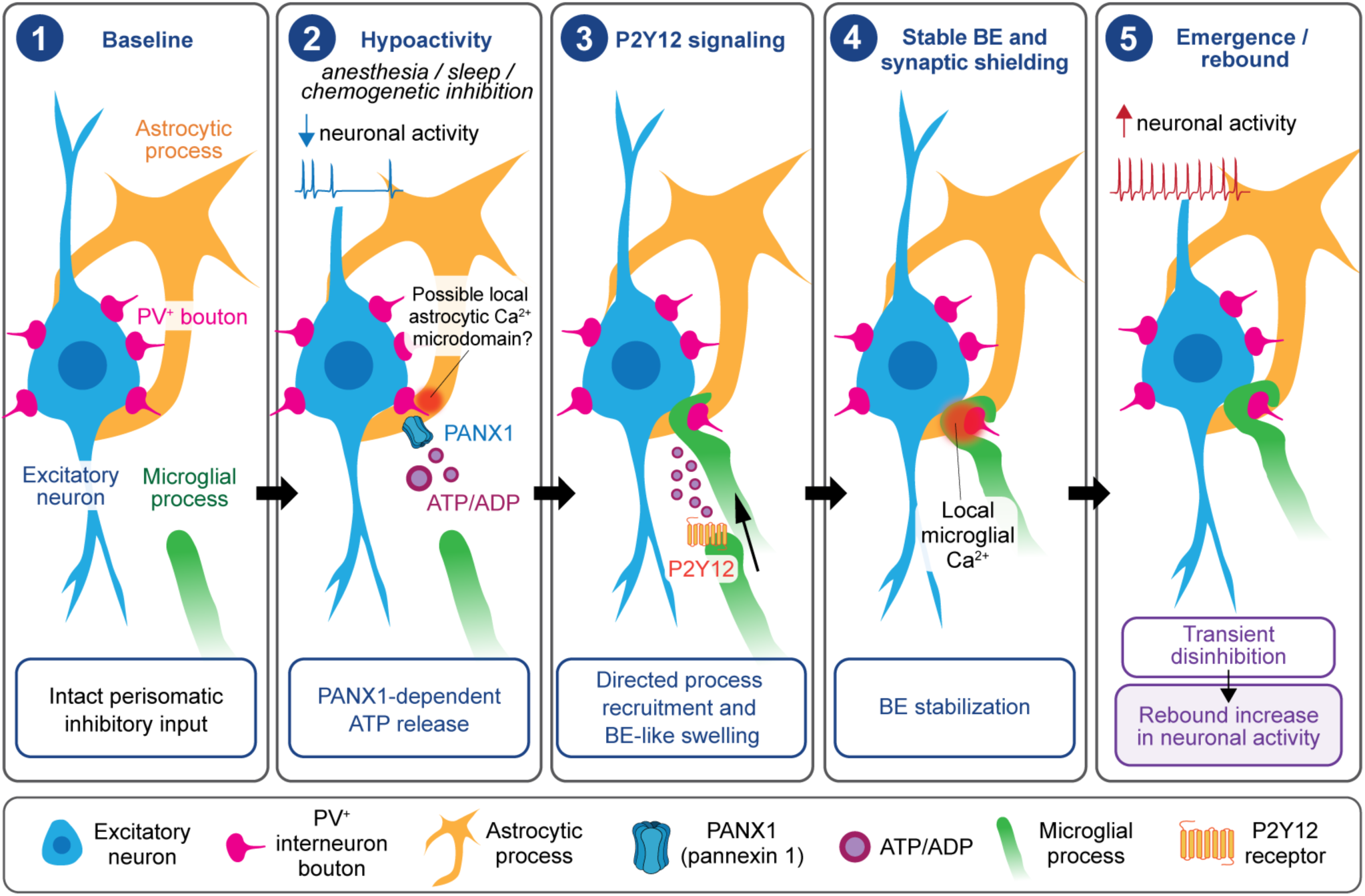
Mechanistic flow of hypoactivity-induced astrocyte-to-microglia purinergic signaling in synaptic shielding. (1) Under baseline conditions, PV⁺ boutons provide perisomatic inhibitory input onto excitatory neuronal somata, while microglial processes survey the surrounding parenchyma. (2) During neuronal hypoactivity associated with anesthesia, sleep, or chemogenetic inhibition, reduced neuronal activity and state-dependent neuromodulatory changes, including reduced noradrenergic tone, create a permissive environment for local glial engagement. Astrocytic PANX1-dependent ATP release generates a localized extracellular nucleotide signal, depicted as ATP/ADP. Spatially restricted astrocytic Ca²⁺ events observed near PV boutons during anesthesia may be associated with local ATP release through PANX1. (3) ATP/ADP engages microglial P2Y12 signaling, promoting directed process recruitment and BE-like swelling toward local nucleotide signals. (4) Localized microglial Ca²⁺ activity accompanies the stabilization of bulbous endings (BEs), which form near PV boutons and shield perisomatic inhibitory synapses. (5) During emergence from anesthesia, persistent BE-mediated synaptic shielding is proposed to transiently attenuate inhibitory input, thereby contributing to disinhibition and a rebound increase in neuronal activity. BE, bulbous ending; PANX1, pannexin 1; PV, parvalbumin.

## Supplementary Movies

**Movie S1. Local ATP hotspot dynamics during awake, isoflurane anesthesia, and emergence.** Representative in vivo two-photon time-lapse imaging of GRAB-ATP1.0 fluorescence in layer 2/3 of the motor cortex during isoflurane anesthesia, corresponding to Fig. 1F. A spatially confined, transient increase in ATP sensor fluorescence indicates a local ATP hotspot event.

**Movie S2. Formation of a microglial bulbous ending associated with a large ATP hotspot.** Representative in vivo two-photon time-lapse imaging of GRAB-ATP1.0 fluorescence and Cx3cr1-GFP-positive microglial morphology (green), together with jRGECO1a-labeled excitatory neurons (red), during isoflurane anesthesia, corresponding to Fig. 2B. A microglial process extends toward a large ATP hotspot (≥100 μm²) and forms a persistent bulbous ending at the ATP-associated site. Arrows indicate ATP release sites, and arrowheads indicate bulbous endings.

**Movie S3. Formation of a microglial bulbous ending associated with a small ATP hotspot.** Representative in vivo two-photon time-lapse imaging of GRAB-ATP1.0 fluorescence and Cx3cr1-GFP-positive microglial morphology (green), together with jRGECO1a-labeled excitatory neurons (red), during isoflurane anesthesia, corresponding to Fig. 2B. A microglial process extends toward a small ATP hotspot (<100 μm²) and forms a persistent bulbous ending at the ATP-associated site. Arrows indicate ATP release sites, and arrowheads indicate bulbous endings.

**Movie S4. ATP-associated microglial bulbous-ending formation near a PV bouton.** Representative in vivo two-photon time-lapse imaging of GRAB-ATP1.0 fluorescence and Cx3cr1-GFP-positive microglia (green), together with tdTomato-labeled PV boutons (magenta), during isoflurane anesthesia, corresponding to Fig. 2I. A local ATP hotspot is followed by microglial process recruitment and bulbous-ending formation near a PV bouton. Arrows indicate ATP release sites, and arrowheads indicate BEs associated with PV boutons.

**Movie S5. Localized astrocytic Ca²⁺ activity near PV boutons during isoflurane anesthesia.** Representative in vivo two-photon time-lapse imaging of GCaMP6f-reported astrocytic Ca²⁺ activity (green) and tdTomato-labeled PV boutons (magenta) across awake, isoflurane anesthesia, and emergence, corresponding to Fig. 3L. Overall Ca²⁺ activity in astrocytes is reduced, whereas spatially confined astrocytic Ca²⁺ events persist near PV boutons during anesthesia. Arrowheads indicate localized astrocytic Ca²⁺ events.

**Movie S6. Localized microglial Ca²⁺ activity within a bulbous ending in a P2Y12 WT mouse.** Representative in vivo two-photon time-lapse imaging of microglial Ca²⁺ activity (green) and microglial morphology (magenta) in a P2Y12 WT Cx3cr1-GCaMP7s-tdTomato mouse, corresponding to Fig. 4E. GCaMP7s fluorescence reveals compartmentalized Ca²⁺ activity within a microglial bulbous ending.

**Movie S7. ATP-associated bulbous-ending formation and localized microglial Ca²⁺ activity in a P2Y12 WT mouse.** Representative simultaneous in vivo two-photon time-lapse imaging of extracellular ATP dynamics and microglial Ca²⁺ activity (green), together with microglial morphology (magenta), in a P2Y12 WT mouse during isoflurane anesthesia, corresponding to Fig. 4G. A local ATP event is followed by bulbous-ending formation and localized Ca²⁺ activity within the bulbous ending. Time is shown as HH:MM:SS, with 00:00:00 indicating the onset of ATP release.

**Movie S8. Transient ATP-associated process-tip swelling in a P2Y12 KO mouse.** Representative simultaneous in vivo two-photon time-lapse imaging of extracellular ATP dynamics and microglial Ca²⁺ activity (green), together with microglial morphology (magenta), in a P2Y12 KO mouse during isoflurane anesthesia, corresponding to Fig. 4G. A local ATP event is followed by transient process-tip swelling and subsequent retraction, without detectable localized microglial Ca²⁺ activity. Time is shown as HH:MM:SS, with 00:00:00 indicating the onset of ATP release.

**Movie S9. Local ATP release and microglial bulbous-ending formation following chemogenetic neuronal silencing.** Representative in vivo two-photon time-lapse imaging of GRAB-ATP1.0 fluorescence and Cx3cr1-GFP-positive microglial morphology (green), together with mCherry-labeled Gi-DREADD-expressing excitatory neurons (magenta), following DCZ administration, corresponding to Fig. 5B. The circle denotes the soma of a DREADD-expressing neuron. A local ATP hotspot event is followed by microglial process recruitment and bulbous-ending formation near Gi-DREADD-expressing neurons. Time is shown as HH:MM:SS, with 00:00:00 indicating the onset of ATP release.

